# The effects of an 8-week mindful eating intervention on anticipatory reward responses in striatum and midbrain

**DOI:** 10.1101/165845

**Authors:** Lieneke K Janssen, Iris Duif, Anne EM Speckens, Ilke van Loon, Joost Wegman, Jeanne HM de Vries, Roshan Cools, Esther Aarts

## Abstract

Obesity is a highly prevalent disease, usually resulting from chronic overeating. Accumulating evidence suggests that increased neural responses during the anticipation of high-calorie food play an important role in overeating. A promising method for counteracting enhanced food anticipation in overeating might be mindfulness-based interventions (MBIs). However, the neural mechanisms by which MBIs can affect food reward anticipation are unclear. In this randomized, actively controlled study, the primary objective was to investigate the effect of an 8-week mindful eating intervention on reward anticipation. On the neural level, we hypothesized that mindful eating would decrease striatal reward anticipation responses. Additionally, responses in the midbrain – from which the reward pathways originate – were explored. Using functional magnetic resonance imaging (fMRI), we tested 58 healthy participants with a wide body mass index range (BMI: 19-35 kg/m^2^), motivated to change their eating behavior. During scanning they performed an incentive delay task, measuring neural reward anticipation responses to caloric and monetary cues before and after 8 weeks of mindful eating or educational cooking (active control). Compared with the educational cooking intervention, mindful eating affected neural reward anticipation responses, with relatively reduced caloric versus monetary reward responses. This effect was, however, not seen in the striatum, but only in the midbrain. The secondary objective was to assess temporary and long-lasting (one year follow-up) intervention effects on self-reported eating behavior and anthropometric measures (BMI, waist circumference, waist-to-hip-ratio (WHR)). We did not observe effects of the mindful eating intervention on eating behavior. Instead, the control intervention showed temporary beneficial effects on BMI, waist circumference, and diet quality, but not on WHR or self-reported eating behavior, as well as long-lasting increases in knowledge about healthy eating. These results suggest that an 8-week mindful eating intervention may have decreased the relative salience of food cues by affecting midbrain but not striatal reward responses. However, these exploratory results should be verified in confirmatory research.

The primary and secondary objectives of the study were registered in the Dutch Trial Register (NTR): NL4923 (NTR5025).

## Introduction

Reward-related disorders such as addiction, binge-eating disorder and obesity, are characterized by altered responses to reward cues related to the target of abuse ^1–3^. Mesolimbic regions in the brain, including the striatum and the midbrain – with its dopaminergic projections to the striatum^4,5^ – respond to increases in appetitive motivation induced by reward cues ^6^. Responses of these subcortical reward regions have been related to eating behavior. For example, greater ventral striatal responses to reward cues have been associated with subsequent food intake ^7^ and future weight gain ^7–9^ (for a review, see ^3^). Reductions in striatal food-cue responses after a weight loss intervention were even predictive of the later outcome of the weight loss intervention^10^. Moreover, increases in BMI were associated with increased midbrain responses to high-calorie food cues in adults ^11^ and to anticipating rewards during risky choices in adolescents ^12^. Interventions targeted at diminishing subcortical responses to food reward cues may therefore be promising for treating and preventing obesity.

Mindfulness-based interventions are aimed at cultivating attention to present-moment experience, without judgment ^13^. Protocolized mindfulness interventions, such as mindfulness-based stress reduction (MBSR) have shown to be effective in reducing subcortical responses to emotional stimuli in anxiety ^14^ as well as in healthy individuals ^15^. Furthermore, mindfulness meditation training can improve executive control processes such as conflict monitoring and response inhibition ^16^, as well as alter functional connectivity of brain networks involved in attention, cognitive processing, awareness, sensory integration, and reward processing ^17^.

Importantly, mindfulness-based interventions aimed at changing eating behavior were able to reduce obesity-related eating behavior in clinical populations ^18,19^, as well as abdominal fat ^20,21^, and to increase self-reported mindful eating ^22^ and reduce reward-driven eating in obese individuals ^23^. However, only two of these trials were actively controlled ^18,22,23^. It is therefore unclear whether these beneficial effects can be attributed to mindfulness per se. In fact, Kristeller and colleagues ^18^ found that both mindfulness-based eating awareness training (MB-EAT) and a psycho-educational/cognitive-behavioral (i.e., active control) intervention decreased binge-eating symptoms relative to a waitlist control group to a similar degree. Given the different nature of these interventions, it is possible that reduced symptomatology was mediated by distinct brain mechanisms, as was suggested by an actively controlled clinical trial on social anxiety ^14^. In this fMRI study, reduced social anxiety symptoms were observed for both the mindfulness and the active control intervention, but the interventions had differential effects on neural responses during self-referential processing. Studies investigating the neurocognitive mechanism underlying mindful eating are required to assess whether a mindful eating intervention can diminish neural responses to food reward cues.

Kirk and colleagues performed three studies on neurocognitive reward mechanisms underlying mindfulness. They found that meditators, relative to controls, showed lower neural responses in striatum during reward anticipation ^24^, as well as diminished BOLD responses in putamen during positive and negative prediction errors ^25^. In addition, they found that mindfulness training modulated value signals in vmPFC to primary reward (juice) delivery ^26^. However, these studies do not yet address the question how mindfulness training affects neural responses for food reward anticipation. Specifically, the first two studies were performed in experienced meditators versus controls instead of in a randomized controlled design, and the third study investigated reward responses at the moment of reward delivery, instead of anticipation. Reward anticipation is particularly interesting to investigate in light of overweight and obesity, as increases in reward anticipation have predictive value for weight gain or overeating-related behavior in these disorders ^1–3,7–9,12^.

Here, we present an actively controlled randomized study investigating the effects of mindfulness on reward anticipation in the brain. We studied the effects of an 8-week mindful eating intervention aimed at changing undesired eating habits versus a carefully matched educational cooking intervention (active control). To assess reward anticipation, we used an incentive delay task ^27^ during fMRI, which has been shown to produce reliable mesolimbic responses to reward cues ^5^. We hypothesized that the mindful eating intervention would reduce reward cue responses in the striatum (primary objective), and also explored these effects in the dopaminergic midbrain as part of the mesolimbic reward circuit. We included both monetary and caloric rewards in the task, which enabled us to assess whether the effect on anticipatory reward responses is specific to the caloric domain, or generalizes to the monetary domain. As a secondary objective, we assessed the effects of mindful eating on anthropometric measures (BMI, waist-to-hip ratio (WHR), and waist circumference) and on self-reported questionnaires related to eating behavior and knowledge of healthy eating.

## Materials and methods

### Participants

The results reported in this study are based on data from 58 healthy, right-handed participants (48 women; mean age: 31.6, SD: 11.0, range: 19 – 52 years; mean body mass index (BMI): 26.0, SD: 3.68, range: 19.7 – 34.7 kg/m2). Note that this sample is largely overlapping with the sample reported previously for another task ^28^. Participants were recruited from Nijmegen and surroundings through advertisement. Only participants (aged: 18 – 55 years old; BMI: 19 – 35 kg/m^2^) with no (history of) eating disorders or current dieting and who were highly motivated to change their eating behavior (not to lose weight per se) were included in the study.

Exclusion criteria included MRI-incompatibility; hepatic, cardiac, respiratory, renal, cerebro-vascular, endocrine, metabolic, pulmonary, or cardiovascular diseases; eating, neurological, or psychiatric disorders; use of neuroleptica or other psychotropic medication; sensori-motor handicaps; drug or alcohol addiction; current strict dieting and a change in body weight of more than 5 kg in the past two months. Crucially, subjects with previous MBSR (Mindfulness-Based Stress Reduction) or MBCT (Mindfulness-Based Cognitive Therapy) experience were excluded from the study. Exclusion criteria are further detailed in Janssen et al. ^28^.

Ten participants were excluded from the analyses following testing because of technical problems (n=6), excessive movement during fMRI scanning (n=1), an incidental finding after the post-test session (n=1), or because of poor task performance (n=2) (for details see **Methods, Behavioral analyses**). For a flow diagram of all excluded participants, see **Figure 1**.

**Figure 1.**
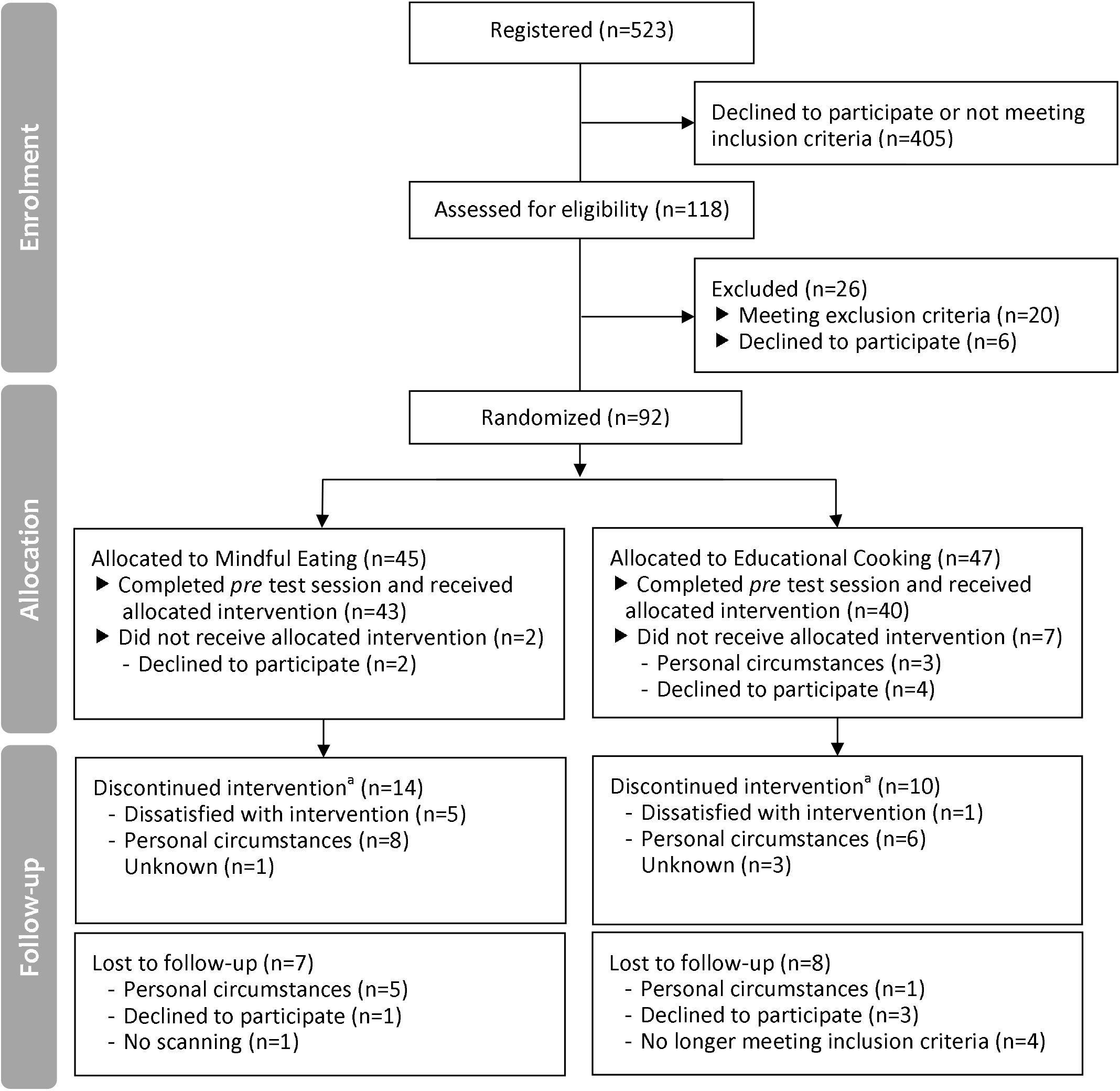
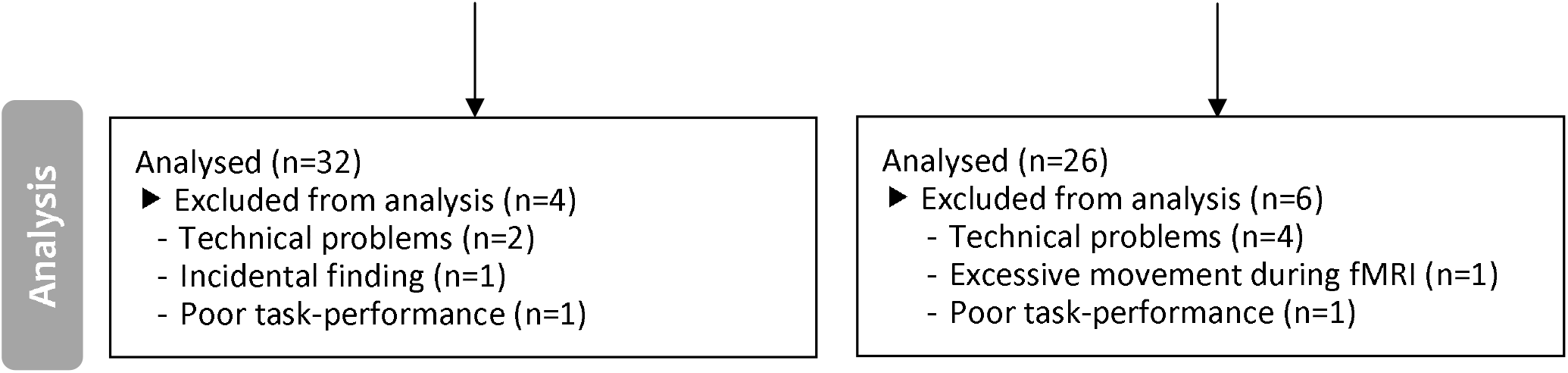
CONSORT flow diagram. ^a^ Attended <4 sessions of the intervention program. Note that these participants were invited back to the laboratory for the post-intervention test session.

All participants gave written informed consent and were reimbursed for participation according to the local institutional guidelines (i.e., 8 Euros per hour for behavioral testing, 10 Euros per hour for scanning). The study protocol was approved by the local ethics committee (CMO region Arnhem-Nijmegen, the Netherlands, 2013-188) and was in accordance with the Declaration of Helsinki. The trial was registered at the Dutch trial register (NL4923 (NTR5025)).

### Protocol

In a separate screening interview, all participants were assessed for in- and exclusion criteria, and matching criteria (age, gender, BMI, experience with meditation and yoga) by taking anthropometric measures and administering self-report questionnaires.

After inclusion, participants came to the MRI laboratory twice – before and after the intervention – and a third time to the behavioral lab one year later. Participants were instructed to abstain from eating foods and drinking anything else than water four hours prior to the start of the test sessions. Participants were also instructed to abstain from drinking alcohol 24 hours before the test session. As secondary outcome measures, anthropometric measurements were taken (weight, height, waist and hip circumference) before scanning and participants completed self-reported measures of diet quality and eating behaviour: the Dutch Healthy Diet - Food Frequency Questionnaire ^29^ (DHD-FFQ) on food intake; a shortened version of the Food Behavior Questionnaire (FBQ) with subscales on “knowledge of healthy eating” and “temptation”; and the Dutch Eating Behaviour Questionnaire ^30^ (DEBQ) with subscales on restraint, emotional, and external eating behaviors. To further characterize the sample, to account for between-group differences at baseline that could occur by chance, and to further explore the effectiveness of the intervention programs, the following self-report questionnaires and scales were administered: the Five Facet Mindfulness Questionnaire – Short Form ^31^ (FFMQ-SF); a Treatment Credibility Questionnaire (TCQ); the Positive And Negative Affect Scale ^32^ (PANAS); the Behavioral Inhibition System / Behavioral Approach System questionnaire ^33^ (BIS-BAS); the Hospital Anxiety and Depression Scale ^34^ (HADS); the Fagerstrom Test for Nicotine Dependence ^35^ (FTND); the Barratt Impulsiveness Scale-11 ^36^ (BIS-11); the Kirby monetary choice, delay discounting questionnaire ^37^; and the neuropsychological digit span test ^38^. Note that the pre-training TCQ was filled out at the first training session, not on the pre-training test session, as participants were unaware of the contents of their training at that time.

After completing the questionnaires, participants underwent a one-hour MR scanning session in which they performed an incentive delay task. Participants also performed a food Stroop task inside the scanner, followed by a reversal learning and outcome devaluation task outside the scanner. These data are reported elsewhere ^28,39,40^. One year after the intervention, participants were re-invited to the laboratory to reassess anthropometric measurements of obesity (weight, waist and hip circumference) and the self-report questionnaires as administered on pre- and post-test sessions. Reward anticipation was not re-assessed at one-year follow-up. The procedure is further detailed in ^28^.

### Paradigm: Incentive Delay task

We adapted the original incentive delay task ^27^ to assess reward anticipation following monetary as well as caloric cues. For task details, see **Figure 2**. In short, on each trial participants were cued as to which of four rewards they could win (monetary: 1 or 50 cents; caloric: a sip of water or of a high-calorie drink of their choice (orange juice, whole chocolate milk or regular cola)). As soon as a white star (target) appeared on the screen, participants were to press a button with their right index finger as fast as possible. If participants responded within an individually determined time-window, they won and the reward was added to their cumulative gain. On average, 59.6% (SD: 10.0) of the trials were hit trials. After scanning, participants received and drank their total caloric gain. Their total monetary gain was added to their financial reimbursement. Participants received instructions for the incentive delay task before going into the scanner, and were aware they would receive their gain following scanning. Before scanning, participants rated how much they *wanted* and *liked* each reward on a Visual Analogue Scale (VAS, 100mm). To expose participants to the reward outcomes, they were provided with the actual coins, and one sip (5 mL) of water and one of the chosen drink while rating the VAS.

**Figure 2.**
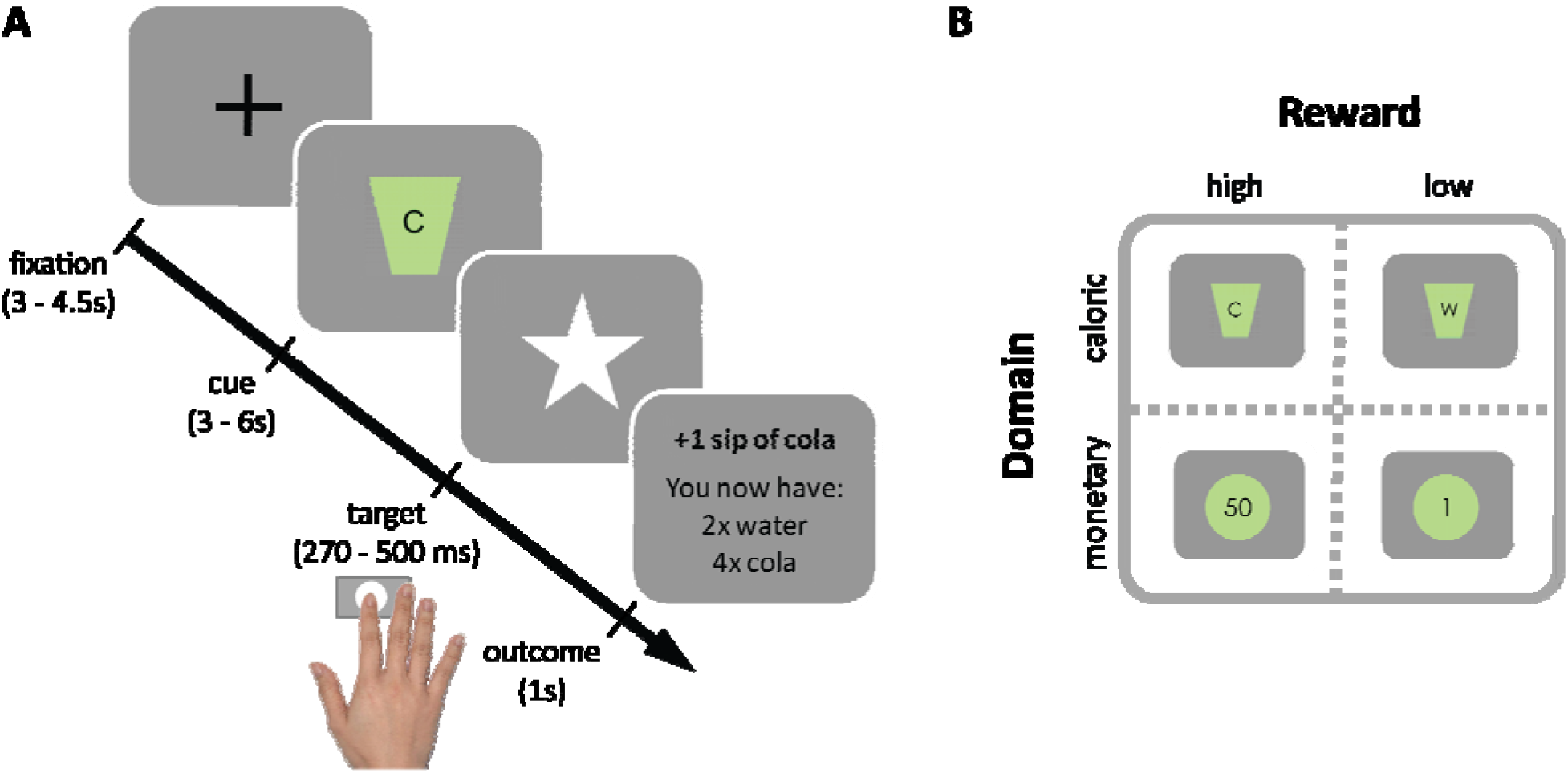
Incentive delay task. A) Each trial started with a fixation cross, followed by a cue signaling which reward could be earned on that trial. Subsequently, a white star (i.e. target) appeared for a brief period and participants were instructed to press a button as fast as possible upon detection using their right index finger. If participants pressed before the response deadline (hit trial), the target remained on the screen, informing participants of the successful registration of their key press. Subsequently, a brief feedback image informing the participants about the total gain was presented. If participants pressed too late or failed to press at all (too late or miss trial, respectively), they were presented with the text message “you win nothing” plus the total gain so far. To ensure participants won similar amounts of each reward (in ±2/3 of the trials), target presentation times were determined individually and adaptively: following hit trials the response deadline for that reward cue was decreased with 10 ms, following too late or miss trials it increased with 10 ms. B) Reward cues for high- and low-calorie cues (C: participant’s choice from cola, orange juice or chocolate milk vs. W: water) and high and low monetary cues (50 cents vs. 1 cent). The task took between 20 – 25 minutes to complete. Participants performed 4 blocks of 25 trials (a total of a 100 trials). A block contained either high/low monetary or high/low-calorie trials. Each trial type was repeated approximately 25 times (M: 24.4, SD: 2.78). Block-presentation was pseudo-randomly distributed and counterbalanced across participants (randomization scheme: ABBA or BAAB).

### Interventions

Participants were randomly assigned to one of two intervention programs: mindful eating (ME) or educational cooking (EC; active control). Participants were assigned by a computer through minimization ^41^, which guarantees that groups are balanced in terms of certain *a priori* determined minimization factors: age (categories: 18-25y, 26-35y, 36-45y, 46-55y), gender (categories: male, female), BMI (categories: 19 – 24.9 kg/m^2^ normal weight, 25 – 29.9 kg/m^2^ overweight, 30 – 35 kg/m^2^ moderately obese) and experience with meditation and yoga (categories: never, 0 – 2 years, 2 – 5 years, 5 – 10 years, > 10 years).

The intervention programs were matched in terms of time, effort, and group contact, but differed significantly in terms of content. Both programs consisted of 8 weekly, 2.5 hour group sessions plus one day (6 hours) dedicated to the intervention goals. Participants were asked to spend 45 minutes per day on homework assignments and to record the amount of time spent on homework forms. In the information letters, the intervention programs were described as “eating with attention” (ME) and “eating with knowledge” (EC) to prevent a selection-bias of participants interested in mindfulness. Only after the first test session, participants were informed about the intervention to which they were randomized, to ensure that baseline measurements were not influenced by intervention expectations. Because group size was set to 10 to 15 participants per round, included participants were divided across three rounds for each intervention (3xME, 3xEC). The final sample for statistical analyses consisted of 32 (from 45 included) participants in the ME intervention and 26 (from 47 included) participants in the EC intervention (for a flow diagram see **Figure 1**). Despite the numerical difference in dropouts between groups, the number of people excluded from analysis was not significantly different (ME: 28.8%, EC: 44.7%, χ^2^(1, N = 92) = 2.461, *ρ* = .117). We get back to the relatively high dropout rate in the **Discussion**.

#### Mindful eating (ME)

The aim of the ME intervention was to increase experiential awareness of food and eating. The ME program was based on the original MBSR program developed by Kabat-Zinn et al. ^42^. Participants performed formal mindfulness practices (i.e. body scan, sitting meditation, walking meditation and mindful movement), aimed at increasing general mindfulness skills, which were similar to the original program. In addition, participants performed informal mindfulness practices based on the Mindful Eating, Conscious Living program (MECL) ^43^, which were mainly directed to mindful eating and not part of the original MBSR program. Sessions focused on themes, such as: the automatic pilot, perception of hunger and satiation, creating awareness of boundaries in eating behavior, stress-related eating, coping with stress, coping with (negative) thoughts, self-compassion, and how to incorporate mindfulness in daily life. Towards the end of the program, participants had a ‘silent day’. During this day, the whole group performed formal mindfulness exercises and ate a meal together in complete silence. Homework consisted of a formal mindfulness practice and an informal mindfulness practice directed at one moment (e.g. a meal) a day. The ME intervention was developed and delivered by qualified mindfulness teachers from the Radboud University Medical Centre for Mindfulness.

#### Educational Cooking (EC)

The aim of the EC intervention was to increase informational awareness of healthy food and eating. The EC program was based on the Dutch healthy food-based dietary guidelines (www.voedingscentrum.nl). To establish similar (active) group activities as in the ME, participants were enrolled in cooking workshops during the group meetings of the EC. Sessions focused on healthy eating, healthy cooking of vegetables and fruit, use of different types of fat and salt for cooking, reading of nutrition labels on food products, healthy snacking, guidelines for making healthy choices when eating in restaurants, and how to incorporate healthy eating and cooking in daily life. Towards the end of the program, participants had a ‘balance day’, during which the participants adhered to all nutritional health guidelines for every snack and meal. Homework assignments entailed practicing cooking techniques, or grocery shopping with informational awareness, and counting the amount of calorie intake for one meal a day. The EC intervention was developed and delivered by a qualified dietitian from Wageningen University and a professional chef of the Nutrition and Dietetics faculty of the University of Applied Sciences of Arnhem-Nijmegen guided the cooking sessions. The interventions are further detailed in ^28^.

### Behavioral analyses

Between-group comparisons were analyzed using independent-samples t-tests, Fisher’s Exact Tests, or Mann-Whitney U tests. Effects of training on anthropometric, neuropsychological and self-report measurements were analyzed using repeated-measures ANOVA with Time (pre, post) as within-participant factor and Intervention (ME, EC) as between-participant factor. To assess the longevity of measures that exhibited a significant Time x Intervention interaction, we ran *post hoc* ANOVAs adding the one-year follow-up data as a third level in factor Time for BMI, waist, DHD-FFQ, and FBQ knowledge. One-year follow-up data was available of 26 participants in the ME group and 21 participants in the EC group. In case of violation of the assumption of sphericity as indicated by Mauchly’s test, the Huyhn-Feldt correction was used to adjust the degrees of freedom accordingly (see Results). Planned *post hoc* comparisons were performed to statistically compare follow-up data to data from both the pre- and post-test sessions separately. Mean latencies of the manual responses were analyzed using repeated-measures ANOVA with within-participant factors Reward (high, low), Domain (caloric, monetary), Time, and the between-participant factor Intervention (ME, EC). Specific effects were tested with subsequent F-tests. All analyses were performed using two-tailed tests in SPSS (version 23.0, Chicago, IL). The significance level was set at an alpha of p=0.05, partial eta squared (η_p_^2^) was reported to indicate effect sizes in the repeated measures ANOVAs.

### fMRI acquisition

We acquired whole-brain functional images (multi-echo) on a Siemens 3T Skyra MRI scanner (Siemens Medical system, Erlangen, Germany) using a 32-channel coil to measure blood oxygen level dependent (BOLD) contrast. A multi-echo echo-planar imaging (EPI) sequence was used to acquire 34 axial slices per functional volume in ascending direction (voxel size 3.5×3.5×3mm; repetition time (TR) 2070 ms; TE 9ms, 19.25ms, 29.5ms, and 39.75ms; flip angle 90 ◻; field of view 224mm). This is a method that uses accelerated parallel imaging to reduce image artifacts (in plane acceleration 3) and acquire images at multiple TEs following a single excitation ^44^. Before the acquisition of functional images, a high-resolution anatomical scan was acquired (T1-weighted MPRAGE, voxel size 1×1×1mm, TR 2300ms, TE 3.03ms, 192 sagittal slices, flip angle 8 ◻, field of view 256 mm).

### fMRI pre-processing and analysis

Data were pre-processed and analyzed using FSL version 5.0.11, (http://www.fmrib.ox.ac.uk/fsl/) and SPM8 (www.fil.ion.ucl.ac.uk/spm). Pre-processing and data analysis were performed using three approaches, which differed in how motion-related noise was accounted for. The final approach was determined based on the strength of the main task effect (i.e. the t-value of the high>low reward anticipation contrast) independent of training, across all participants and sessions. First, we added twelve rigid-body transformation parameters (three translations and rotations, and their linear derivatives) obtained during realignment to the first level model. Second, we used non-aggressive ICA-AROMA ^45^ to reduce motion-induced signal variations in the fMRI data. Because ICA-AROMA takes out noise components, the twelve rigid-body transformation parameters obtained during realignment were not included in the model. For our third approach, we also used ICA-AROMA, however, rather than reducing motion-related noise in the fMRI data directly, we added the time courses of the independent components accounting for less than 5% of task-related variance to the first level model. To achieve this, we used the components identified as motion by ICA-AROMA in a multiple regression analysis with the task regressors as predictors and the motion-related time courses as dependent variables. From this analysis, the adjusted R^2^ was obtained to identify how much of the total variance in a time course was captured by the task’s design. In case the adjusted R^2^ of a component was higher than 5%, they were not included in the first level model as noise regressors (i.e. regressor of non-interest). The twelve rigid-body transformation parameters obtained during realignment were also included in the model. The third approach showed the strongest main task effect (brain responses to high - low reward cues) and was therefore used as our final approach. Below, we describe this approach in more detail.

The volumes for each echo time were realigned to correct for motion artefacts (estimation of the realignment parameters is done for the first echo and then copied to the other echoes). The four echo images were combined into a single MR volume based on 31 volumes acquired before the actual experiment started using an optimized echo weighting method ^44^. Combined functional images were slice-time corrected by realigning the time-series for each voxel temporally to acquisition of the middle slice. The images were subsequently spatially smoothed using an isotropic 6 mm full-width at half-maximum Gaussian kernel. Non-aggressive ICA-AROMA ^45^ was used to identify motion-induced signal variations in the fMRI data. Participant-specific structural and functional data were then coregistered to a standard structural or functional stereotactic space respectively (Montreal Neurological Institute (MNI) template). After segmentation of the structural images using a unified segmentation approach, structural images were spatially coregistered to the mean of the functional images. The resulting transformation matrix of the segmentation step was then used to normalize the anatomical and functional images into Montreal Neurological Institute space. The functional images were resampled at voxel size 2 x 2 x 2 mm.

Statistical analyses of fMRI data at the individual participant (first) level were performed using an event-related approach and included 13 regressors of interest: four regressors for cue presentation (high- and low-calorie cues, high and low monetary cues), one regressor for target presentation, four outcome regressors for hits (high- and low-calorie hits, high and low monetary hits), and four outcome regressors for trials on which participants responded too late (high- and low-calorie too late, high and low monetary too late). If participants failed to respond on a trial (i.e. a miss), the trial was excluded from analyses. Onsets of the regressors were modeled as a stick function (duration=0s) convolved with a canonical hemodynamic response function ^46^. Furthermore, we only added time courses of the independent noise components that accounted for less than 5% of task-related variance to the first level model as regressors of non-interest. Note that the number of these regressors varied per subject and session. In addition, twelve rigid-body parameters, a constant term, and two regressors that reflected signal variation in white matter and cerebrospinal fluid regions were included as regressors of non-interest. High pass filtering (128 seconds) was applied to the time series of the functional images to remove low-frequency drifts and correction for serial correlations was done using an autoregressive AR(1) model.

We ran two general linear models (GLMs) at the second level: one for reward anticipation with high minus low reward cue contrast images, and one for reward receipt with hit minus too late contrast images. Analysis of variance (ANOVA) was performed in a full-factorial design, with between-subject factor Intervention and within-subject factors Time and Domain, resulting in 8 cells. Effects were considered statistically significant when reaching a threshold of p<0.05, family wise error (FWE) corrected for multiple comparisons at the peak level, whole brain or in the *a priori* defined regions of interest (see below). We report whole-brain and small volume corrected (pFWE<.05) effects in **Table 3 and 4**, and show the statistical maps at p<.001 and p<.005 uncorrected thresholds in **Figure 3** for exploratory purposes.

**Figure 3.**
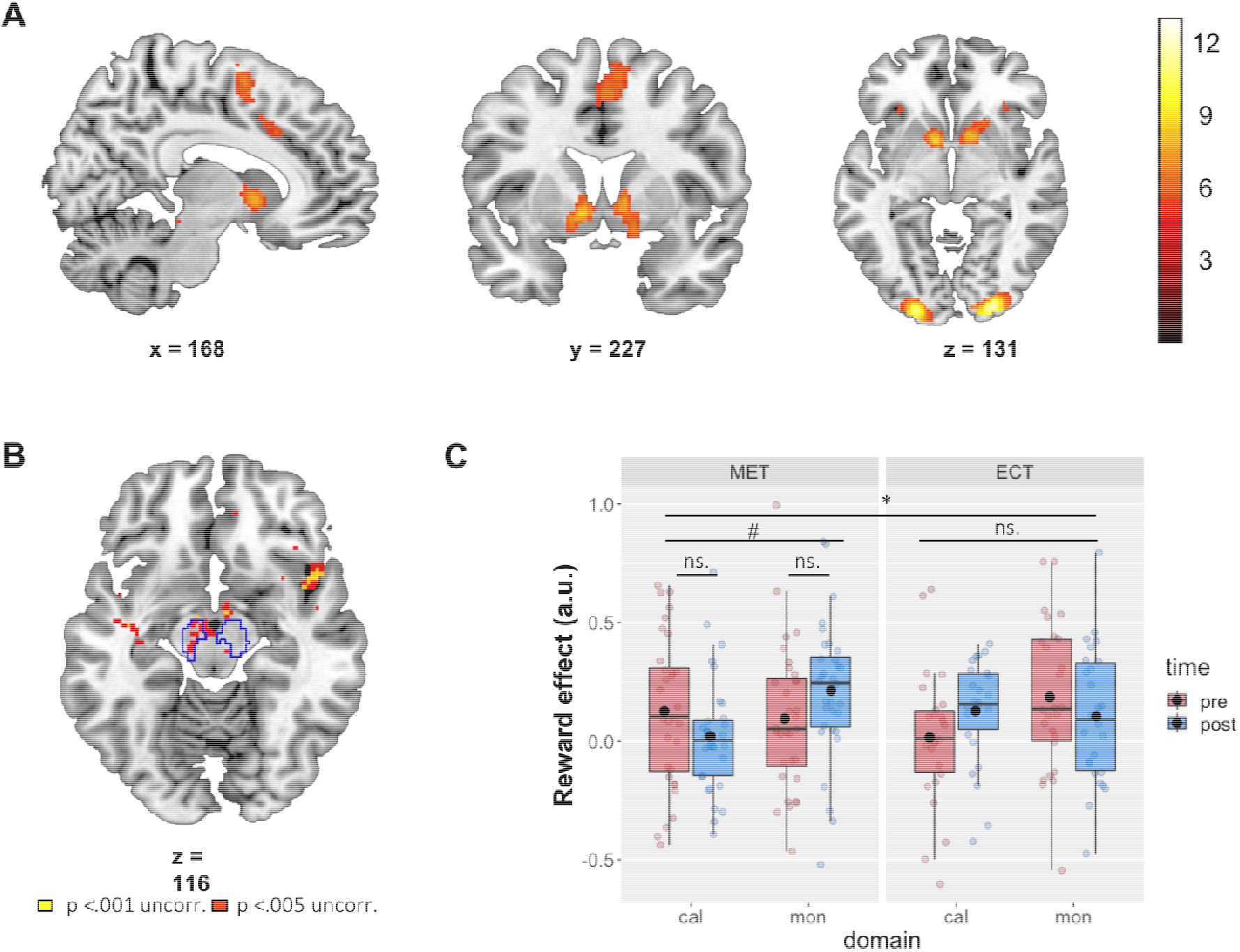
Summary of neuroimaging results. A) Main effect of reward. Contrast of high vs. low reward cue trials (high > low). Full brain statistical parametric maps were thresholded at p < .05 (FWE-corrected, peak-level). B) Axial slice of whole brain interaction effect of Domain x Time x Intervention for the Reward contrast (high > low). Statistical parametric maps were thresholded at p < .001 (yellow) and p < .005 (red) uncorrected for visualization purposes. Outlined regions are corrected for multiple comparisons within our small search volume, at peak pFWE < .05. C) Betas from the bilateral probabilistic midbrain ROI (outlined in blue in panel B). Post-minus pre-intervention mean betas based on the high minus low reward contrast are presented for each domain (caloric, monetary) and for each intervention group (ME, EC) in arbitrary units (a.u.). Box plots show the median and interquartile range, with the black dot denoting the mean. All statistical parametric maps are overlaid onto a T1-weighted canonical image. Slice coordinates are defined in MNI152 space and images are shown in neurological convention (left=left). * = p < .025 (Bonferroni corrected for two ROIs) and # = p < .05.

To further investigate the effects of intervention on reward anticipation and receipt, region-of-interest (ROI) analyses were performed using *a priori* defined ROIs for midbrain and striatum. ROIs were anatomically defined based on a high-resolution probabilistic *in vivo* atlas that included midbrain and striatal nuclei ^47^: bilateral substantia nigra (atlas: region 7), and ventral tegmental area (region 11) for *midbrain*, and bilateral caudate nucleus (region 2), nucleus accumbens (region 3) and putamen (region 1) for *striatum* at 100% overlap. Probabilistically weighted mean beta weights were extracted from all voxels in both ROIs separately using MarsBar ^48^. The probabilistically weighted averaged beta-weights were analyzed per region using ANOVA with the same factors as in the whole-brain analyses. As two ROIs were tested (striatum and midbrain), effects for each total region were considered significant when reaching a threshold of p<.025 (Bonferroni corrected for multiple comparisons). *Post hoc*, the same effects were tested in the striatal sub-regions (bilateral caudate nucleus, nucleus accumbens, and putamen) because striatal sub-regions have been associated with distinct neurocognitive mechanisms.

## Results

### Characterization of intervention groups

The mindful eating (ME) and educational cooking (EC) groups were well matched in terms of the minimization factors age, gender, body mass index (BMI) and experience with meditation and yoga (**Table 1**). Note that the groups tended to differ in terms of educational level. However, *post hoc* correlation analyses revealed no correlations between educational level and the neural effects described below and is therefore unlikely to drive these effects. Furthermore, the total time participants spent on the intervention, and the number of sessions participants attended did not differ significantly between the two groups (**Table 1**).

**Table 1.**
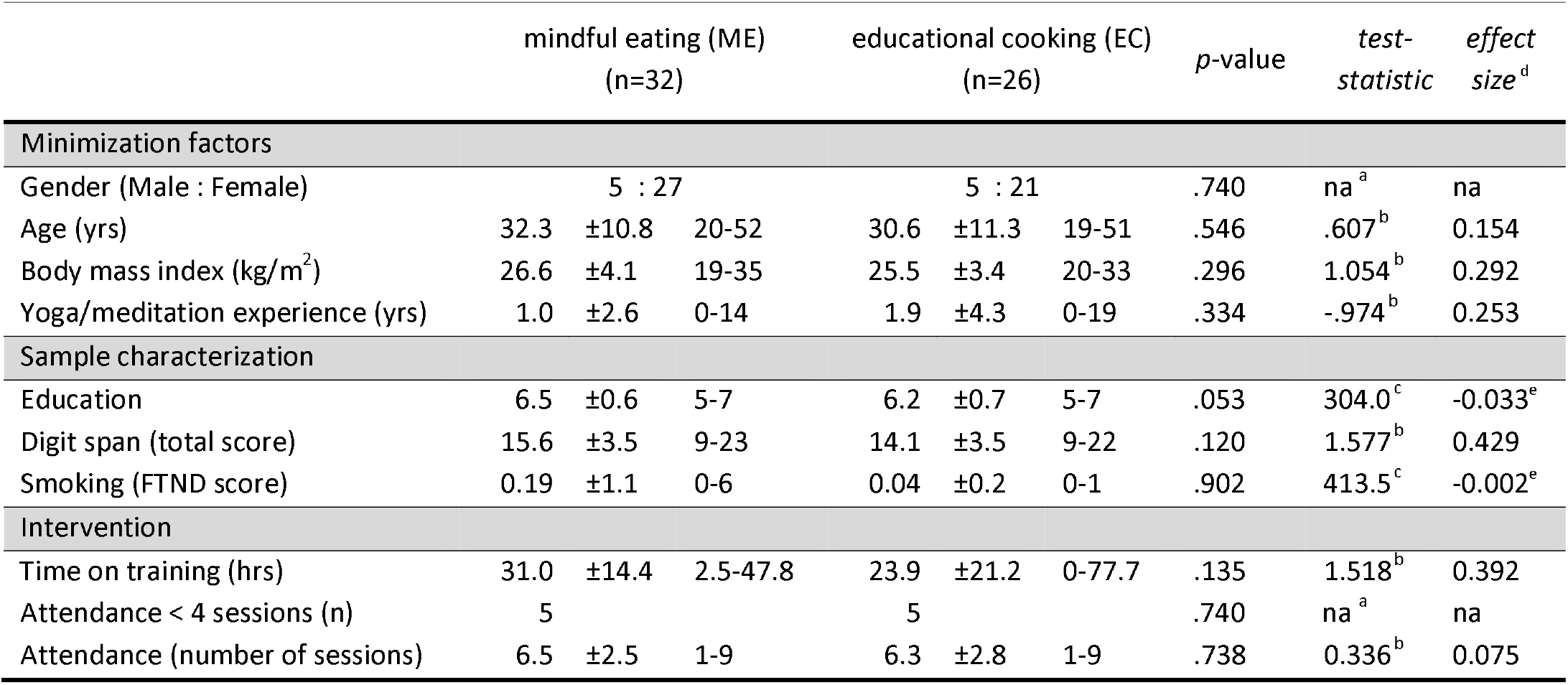
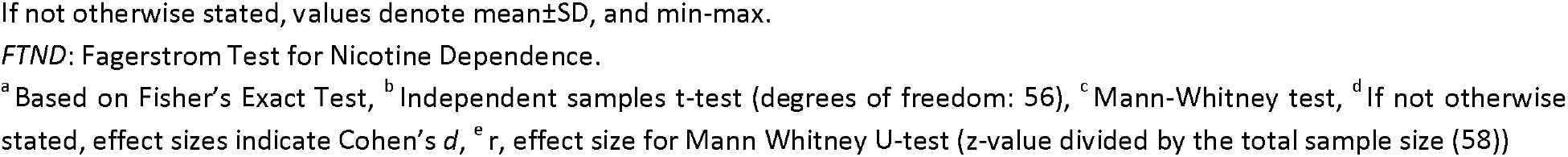
Between-group (mindful eating, ME; educational cooking, EC) comparisons.

|  | mindful eating (ME)<br>(n=32) |  |  | educational cooking (EC)<br>(n=26) |  |  | *p*-value | *test-*<br>*statistic* | *effect*<br>*size<sup>d</sup>* |
| --- | --- | --- | --- | --- | --- | --- | --- | --- | --- |
| Minimization factors |  |  |  |  |  |  |  |  |  |
| Gender (Male : Female) | 5 : 27 |  |  | 5 : 21 |  |  | .740 | na<sup>a</sup> | na |
| Age (yrs) | 32.3 | ±10.8 | 20-52 | 30.6 | ±11.3 | 19-51 | .546 | .607<sup>b</sup> | 0.154 |
| Body mass index (kg/m<sup>2</sup>) | 26.6 | ±4.1 | 19-35 | 25.5 | ±3.4 | 20-33 | .296 | 1.054<sup>b</sup> | 0.292 |
| Yoga/meditation experience (yrs) | 1.0 | ±2.6 | 0-14 | 1.9 | ±4.3 | 0-19 | .334 | -.974<sup>b</sup> | 0.253 |
| Sample characterization |  |  |  |  |  |  |  |  |  |
| Education | 6.5 | ±0.6 | 5-7 | 6.2 | ±0.7 | 5-7 | .053 | 304.0<sup>c</sup> | -0.033<sup>e</sup> |
| Digit span (total score) | 15.6 | ±3.5 | 9-23 | 14.1 | ±3.5 | 9-22 | .120 | 1.577<sup>b</sup> | 0.429 |
| Smoking (FTND score) | 0.19 | ±1.1 | 0-6 | 0.04 | ±0.2 | 0-1 | .902 | 413.5<sup>c</sup> | -0.002<sup>e</sup> |
| Intervention |  |  |  |  |  |  |  |  |  |
| Time on training (hrs) | 31.0 | ±14.4 | 2.5-47.8 | 23.9 | ±21.2 | 0-77.7 | .135 | 1.518<sup>b</sup> | 0.392 |
| Attendance < 4 sessions (n) | 5 |  |  | 5 |  |  | .740 | na<sup>a</sup> | na |
| Attendance (number of sessions) | 6.5 | ±2.5 | 1-9 | 6.3 | ±2.8 | 1-9 | .738 | 0.336<sup>b</sup> | 0.075 |

If not otherwise stated, values denote mean ± SD, and min-max.
*FTND*: Fagerstrom Test for Nicotine Dependence.
<sup>a</sup>Based on Fisher's Exact Test, <sup>b</sup>Independent samples t-test (degrees of freedom: 56), <sup>c</sup>Mann-Whitney test, <sup>d</sup>If not otherwise stated, effect sizes indicate Cohen's *d*, <sup>e</sup>*r*, effect size for Mann Whitney U-test (z-value divided by the total sample size (58))

### Behavioral outcomes

As a primary objective, we assessed the effects of the intervention on reward anticipation during the incentive delay task. We start with the behavioral responses during the task (**Table 2**). Across sessions and intervention groups, participants responded faster on high than on low reward trials (main Reward: F(1,56)=25.0, p < .001, η_p_^2^ = 0.309), thus revealing a reward benefit (**Table 2**). In addition, participants across sessions and intervention groups responded faster to monetary relative to caloric reward cues (main Domain: F(1,56)=17.4, p<.001, η_p_^2^ = 0.237). We observed a reward benefit for both caloric (F(1,56)=4.5, p=.038, η_p_^2^ = 0.074) and monetary trials (F(1,56)=25.6, p<.001, η_p_^2^ = 0.314), which was, however, larger in the monetary trials (Reward x Domain interaction: F(1,56)=9.0, p=.004, η_p_^2^ = 0.139). Participants responded faster on post-relative to pre-intervention test sessions (pre: 310.66 (SD: 21.3), post: 304.60 ms. (SD: 20.8); main Time: F(1,56)=4.4, p<.041, η_p_^2^ = 0.072). However, there was no evidence for effects of intervention type (4-way interaction between Intervention x Time x Reward x Domain (F(1,56)<1), indicating that the speeding of responding on the second versus the first session was not qualified by reward magnitude, reward type or intervention type.

**Table 2.** Task-related outcomes pre- and post-training, for each group (mindful eating, ME; educational cooking, EC) separately, and Time (pre, post) x Intervention (ME, EC) statistics.

|  | mindful eating (ME) |  |  |  | educational cooking (EC) |  |  |  | *p* | *test-*<br>*statistic<sup>a</sup>* | *effect*<br>*size<sup>b</sup>* |
| --- | --- | --- | --- | --- | --- | --- | --- | --- | --- | --- | --- |
|  | pre |  | post |  | pre |  | post |  |  |  |  |
| Primary outcome measure: Response times on the incentive delay task |  |  |  |  |  |  |  |  |  |  |  |
| Response Times per reward type |  |  |  |  |  |  |  |  |  |  |  |
| Low caloric | 313.7 | ±41.0 | 312.4 | ±33.8 | 322.5 | ±51.6 | 312.6 | ±43.8 | .319 | 1.0 | 0.018 |
| High caloric | 303.4 | ±33.8 | 299.1 | ±31.5 | 322.2 | ±50.0 | 311.8 | ±48.4 | .471 | < 1 | 0.009 |
| Low monetary | 313.0 | ±47.0 | 311.2 | ±44.2 | 317.4 | ±44.8 | 313.3 | ±49.6 | .834 | < 1 | 0.001 |
| High monetary | 294.7 | ±26.2 | 285.1 | ±32.5 | 302.3 | ±41.5 | 293.9 | ±43.0 | .874 | < 1 | <0.001 |
| Exploratory outcome measures (manipulation check): Visual analogue scales |  |  |  |  |  |  |  |  |  |  |  |
| Wanting per reward type |  |  |  |  |  |  |  |  |  |  |  |
| Low caloric | 4.5 | ±2.8 | 4.6 | ±2.8 | 4.5 | ±3.1 | 4.6 | ±2.8 | .987 | < 1 | <0.001 |
| High caloric | 6.3 | ±2.0 | 5.8 | ±2.4 | 5.4 | ±3.0 | 5.6 | ±2.4 | .330 | < 1 | 0.017 |
| Low monetary | 1.9 | ±2.4 | 1.5 | ±2.0 | 2.2 | ±2.5 | 2.4 | ±2.6 | .318 | 1.0 | 0.018 |
| High monetary | 5.2 | ±2.8 | 5.4 | ±2.7 | 5.0 | ±3.2 | 5.4 | ±2.4 | .840 | < 1 | 0.001 |
| Liking per reward type |  |  |  |  |  |  |  |  |  |  |  |
| Low caloric | 6.4 | ±2.3 | 6.1 | ±2.2 | 6.2 | ±2.7 | 6.6 | ±2.2 | .187 | 1.8 | 0.031 |
| High caloric | 7.2 | ±1.6 | 6.7 | ±2.1 | 6.8 | ±2.9 | 6.4 | ±2.7 | .783 | < 1 | 0.001 |
| Low monetary | 2.2 | ±2.4 | 2.2 | ±2.2 | 2.8 | ±2.4 | 2.8 | ±2.3 | .967 | < 1 | <0.001 |
| High monetary | 5.1 | ±2.5 | 5.2 | ±2.4 | 4.4 | ±2.7 | 5.3 | ±2.2 | .143 | 2.2 | 0.038 |
| Hunger<sup>c</sup> | 5.9 | ±2.6 | 5.9 | ±2.7 | 5.9 | ±3.0 | 5.6 | ±2.9 | .835 | < 1 | 0.001 |
| Thirst<sup>c</sup> | 5.7 | ±2.6 | 5.9 | ±2.8 | 6.0 | ±2.4 | 5.5 | ±2.4 | .273 | 1.2 | 0.023 |
| Satiety<sup>c</sup> | 2.3 | ±2.1 | 2.1 | ±0.9 | 1.9 | ±1.1 | 2.1 | ±1.2 | .345 | < 1 | 0.017 |

If not otherwise stated, values denote mean±SD.
<sup>a</sup>The reported test-statistic is the F-value (degrees of freedom: 1,56)
<sup>b</sup>The reported effect size is the partial eta squared ( $\eta_p^2$ )
<sup>c</sup>Hunger, Thirst, Satiety: N = 55 (N<sub>ME</sub> = 29, N<sub>EC</sub> = 26; degrees of freedom: 1,53)

There were also no effects of Intervention on any other behavioral task-related measures that we included to control for potentially unexpected group-differences in wanting and liking of the included rewards, or hunger, thirst, and satiety VAS ratings during the task (no Time x Intervention interactions (**Table 2**)).

### Neuroimaging outcomes

#### Reward Anticipation

Before assessing the intervention effects on the neural responses during reward anticipation (primary outcome), we identified brain regions that responded to reward anticipation across sessions and intervention groups (main effect of Reward condition: high>low). At our whole-brain corrected threshold (FWE<.05, peak-level), this contrast yielded significant responses in striatum (right caudate nucleus, right nucleus accumbens, right putamen, and left pallidum) and two right midbrain regions, as well as in occipital, motor and frontal regions (**Figure 3a**). Note that the optimal preprocessing pipeline was selected based on maximal main effects of reward anticipation (see **Materials and Methods**), so no inference can be made on the magnitude of these main effects. Reward anticipation differed in mostly posterior regions for monetary versus caloric reward cues (i.e., interaction of Domain x Reward), independent of sessions and intervention groups. For all contrasts, see **Table 3**.

**Table 3.**
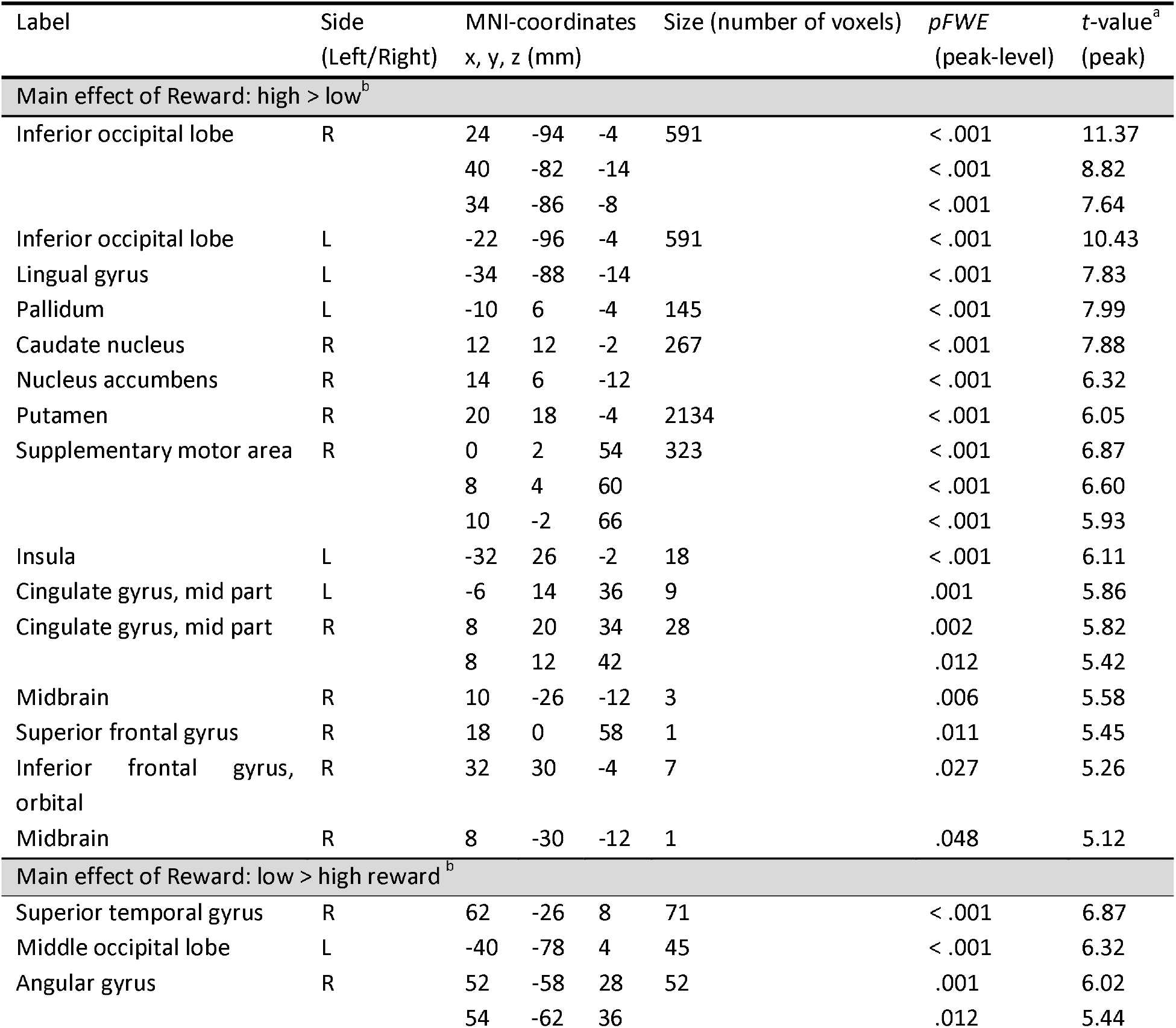
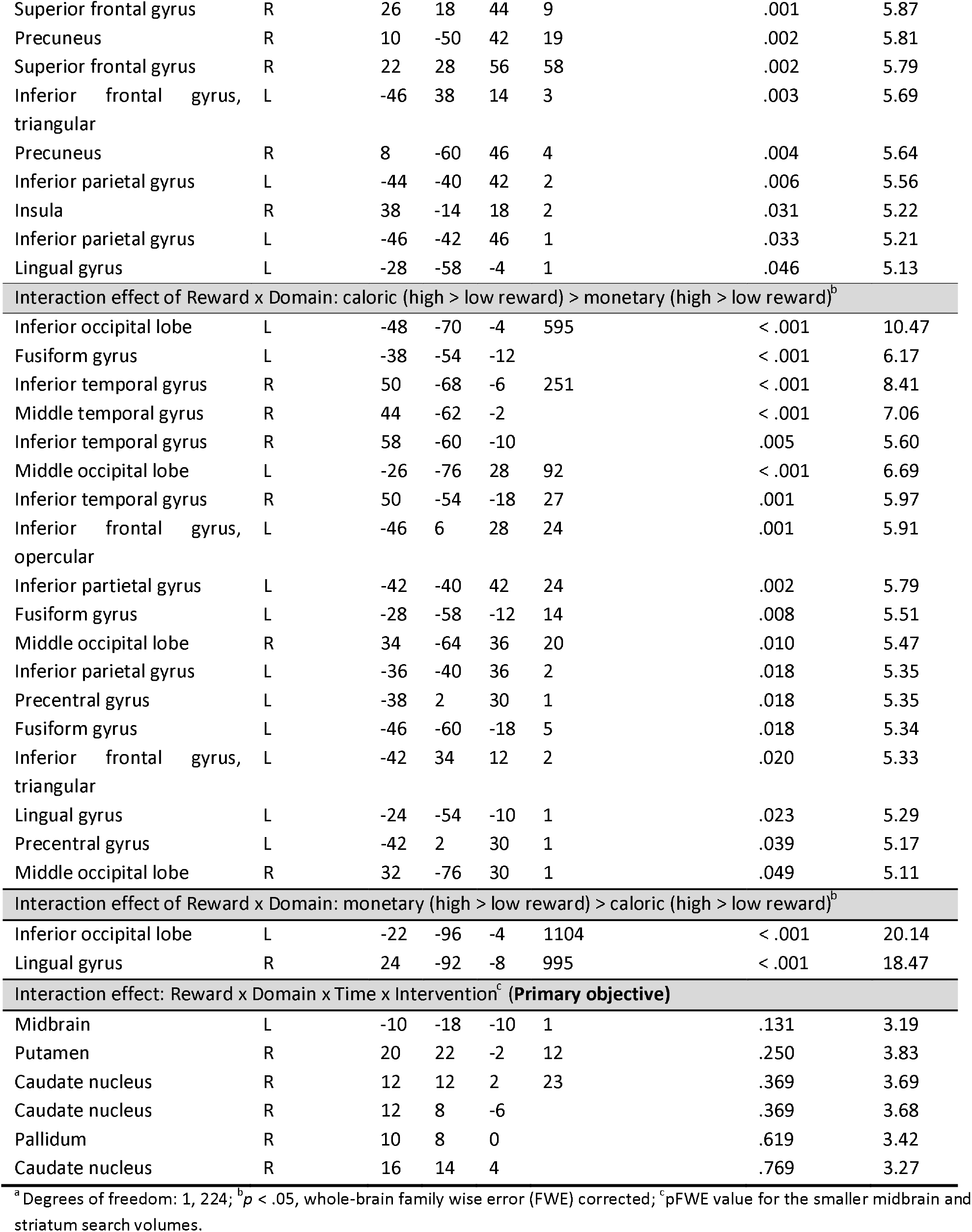
Reward anticipation. Summary of brain regions exhibiting main effects of reward, domain and/or interactions with domain, intervention, and time. N.B., the preprocessing pipeline was selected based on maximal main effects of reward anticipation.

| Label | Side<br>(Left/Right) | MNI-coordinates<br>x, y, z (mm) |  |  | Size (number of voxels) | *p*FWE<br>(peak-level) | *t*-value<sup>a</sup><br>(peak) |
| --- | --- | --- | --- | --- | --- | --- | --- |
| Main effect of Reward: high > low<sup>b</sup> |  |  |  |  |  |  |  |
| Inferior occipital lobe | R | 24 | -94 | -4 | 591 | < .001 | 11.37 |
|  |  | 40 | -82 | -14 |  | < .001 | 8.82 |
|  |  | 34 | -86 | -8 |  | < .001 | 7.64 |
| Inferior occipital lobe | L | -22 | -96 | -4 | 591 | < .001 | 10.43 |
| Lingual gyrus | L | -34 | -88 | -14 |  | < .001 | 7.83 |
| Pallidum | L | -10 | 6 | -4 | 145 | < .001 | 7.99 |
| Caudate nucleus | R | 12 | 12 | -2 | 267 | < .001 | 7.88 |
| Nucleus accumbens | R | 14 | 6 | -12 |  | < .001 | 6.32 |
| Putamen | R | 20 | 18 | -4 | 2134 | < .001 | 6.05 |
| Supplementary motor area | R | 0 | 2 | 54 | 323 | < .001 | 6.87 |
|  |  | 8 | 4 | 60 |  | < .001 | 6.60 |
|  |  | 10 | -2 | 66 |  | < .001 | 5.93 |
| Insula | L | -32 | 26 | -2 | 18 | < .001 | 6.11 |
| Cingulate gyrus, mid part | L | -6 | 14 | 36 | 9 | .001 | 5.86 |
| Cingulate gyrus, mid part | R | 8 | 20 | 34 | 28 | .002 | 5.82 |
|  |  | 8 | 12 | 42 |  | .012 | 5.42 |
| Midbrain | R | 10 | -26 | -12 | 3 | .006 | 5.58 |
| Superior frontal gyrus | R | 18 | 0 | 58 | 1 | .011 | 5.45 |
| Inferior frontal gyrus, orbital | R | 32 | 30 | -4 | 7 | .027 | 5.26 |
| Midbrain | R | 8 | -30 | -12 | 1 | .048 | 5.12 |
| Main effect of Reward: low > high reward<sup>b</sup> |  |  |  |  |  |  |  |
| Superior temporal gyrus | R | 62 | -26 | 8 | 71 | < .001 | 6.87 |
| Middle occipital lobe | L | -40 | -78 | 4 | 45 | < .001 | 6.32 |
| Angular gyrus | R | 52 | -58 | 28 | 52 | .001 | 6.02 |
|  |  | 54 | -62 | 36 |  | .012 | 5.44 |
| Superior frontal gyrus | R | 26 | 18 | 44 | 9 | .001 | 5.87 |
| Precuneus | R | 10 | -50 | 42 | 19 | .002 | 5.81 |
| Superior frontal gyrus | R | 22 | 28 | 56 | 58 | .002 | 5.79 |
| Inferior frontal gyrus, triangular | L | -46 | 38 | 14 | 3 | .003 | 5.69 |
| Precuneus | R | 8 | -60 | 46 | 4 | .004 | 5.64 |
| Inferior parietal gyrus | L | -44 | -40 | 42 | 2 | .006 | 5.56 |
| Insula | R | 38 | -14 | 18 | 2 | .031 | 5.22 |
| Inferior parietal gyrus | L | -46 | -42 | 46 | 1 | .033 | 5.21 |
| Lingual gyrus | L | -28 | -58 | -4 | 1 | .046 | 5.13 |
| Interaction effect of Reward x Domain: caloric (high > low reward) > monetary (high > low reward)<sup>b</sup> |  |  |  |  |  |  |  |
| Inferior occipital lobe | L | -48 | -70 | -4 | 595 | < .001 | 10.47 |
| Fusiform gyrus | L | -38 | -54 | -12 |  | < .001 | 6.17 |
| Inferior temporal gyrus | R | 50 | -68 | -6 | 251 | < .001 | 8.41 |
| Middle temporal gyrus | R | 44 | -62 | -2 |  | < .001 | 7.06 |
| Inferior temporal gyrus | R | 58 | -60 | -10 |  | .005 | 5.60 |
| Middle occipital lobe | L | -26 | -76 | 28 | 92 | < .001 | 6.69 |
| Inferior temporal gyrus | R | 50 | -54 | -18 | 27 | .001 | 5.97 |
| Inferior frontal gyrus, opercular | L | -46 | 6 | 28 | 24 | .001 | 5.91 |
| Inferior parietal gyrus | L | -42 | -40 | 42 | 24 | .002 | 5.79 |
| Fusiform gyrus | L | -28 | -58 | -12 | 14 | .008 | 5.51 |
| Middle occipital lobe | R | 34 | -64 | 36 | 20 | .010 | 5.47 |
| Inferior parietal gyrus | L | -36 | -40 | 36 | 2 | .018 | 5.35 |
| Precentral gyrus | L | -38 | 2 | 30 | 1 | .018 | 5.35 |
| Fusiform gyrus | L | -46 | -60 | -18 | 5 | .018 | 5.34 |
| Inferior frontal gyrus, triangular | L | -42 | 34 | 12 | 2 | .020 | 5.33 |
| Lingual gyrus | L | -24 | -54 | -10 | 1 | .023 | 5.29 |
| Precentral gyrus | L | -42 | 2 | 30 | 1 | .039 | 5.17 |
| Middle occipital lobe | R | 32 | -76 | 30 | 1 | .049 | 5.11 |
| Interaction effect of Reward x Domain: monetary (high > low reward) > caloric (high > low reward)<sup>b</sup> |  |  |  |  |  |  |  |
| Inferior occipital lobe | L | -22 | -96 | -4 | 1104 | < .001 | 20.14 |
| Lingual gyrus | R | 24 | -92 | -8 | 995 | < .001 | 18.47 |
| Interaction effect: Reward x Domain x Time x Intervention<sup>c</sup> (Primary objective) |  |  |  |  |  |  |  |
| Midbrain | L | -10 | -18 | -10 | 1 | .131 | 3.19 |
| Putamen | R | 20 | 22 | -2 | 12 | .250 | 3.83 |
| Caudate nucleus | R | 12 | 12 | 2 | 23 | .369 | 3.69 |
| Caudate nucleus | R | 12 | 8 | -6 |  | .369 | 3.68 |
| Pallidum | R | 10 | 8 | 0 |  | .619 | 3.42 |
| Caudate nucleus | R | 16 | 14 | 4 |  | .769 | 3.27 |

<sup>a</sup> Degrees of freedom: 1, 224; <sup>b</sup> $p < .05$ , whole-brain family wise error (FWE) corrected; <sup>c</sup> pFWE value for the smaller midbrain and striatum search volumes.

We were primarily interested in the effects of ME on reward anticipation in our a priori defined, anatomical region-of-interest (ROI): the striatum. We explored the same effects in an anatomical midbrain ROI. First, we explored these effects using our probabilistic ROIs as small search volumes. We found five peaks for the Reward x Domain x Time x Intervention interaction in the striatum (three regions in caudate nucleus, one in putamen, and one in pallidum), as well as one peak in the midbrain. However, these peaks were not significant when correcting for multiple comparisons across the two search volumes (i.e., midbrain and striatum), i.e., all pFWE > 0.025 (**Figure 3b**).

Based on our hypotheses, we also performed ROI analyses (Figure 3c) using a bilateral probabilistic structural ROI for the striatum (primary) and the midbrain (see Materials and methods). No four-way interaction effect (Intervention x Time x Domain x Reward) was found for the striatum. *Post hoc* analyses of the separate striatal regions also showed no effect of intervention (Intervention x Time x Domain x Reward: putamen: F(1,56)<1, p = .385, η_p_^2^ = 0.014, caudate nucleus: F(1, 56)=1.3, p = .255, η_p_^2^ = 0.023, nucleus accumbens: F(1, 56)=2.2, p = .142, η_p_^2^ = 0.038). Interestingly, for the midbrain ROI, we did observe a significant four-way interaction in the ROI betas, and - in contrast to the observed midbrain effect in the small volume analysis mentioned above - this effect did survive correction for multiple comparisons (Intervention x Time x Domain x Reward: F(1,56)=7.9, p=.007, η_p_^2^ = 0.123, α=.025). Post hoc analyses showed a significant relative reduction in caloric versus monetary reward anticipation in midbrain after the mindful eating training (Time x Domain x Reward for ME: F(1,31)=4.4, p=.043, η_p_^2^ = 0.125). This effect was not significant in the EC group (Time x Domain x Reward for EC: F(1,25)=3.7, p = .065, η_p_^2^ = 0.130) and, if anything, showed the opposite effect. When further breaking down the interaction in the mindfulness group, we found no significant training effect in the caloric domain (Time x Reward: F(1,31)=2.1, p = .156, η_p_^2^ = 0.064), or in the monetary domain (Time x Reward: F(1,31)=2.8, p = .104, η_p_^2^ = 0.083) separately. This means that we can only interpret the ME effect on midbrain reward anticipation responses as a relative decrease for caloric versus monetary reward (see above-mentioned significant Time x Domain x Reward effect for ME). Pre-intervention Reward differences could not explain the observed interaction in the midbrain (caloric: t(56)=1.4, p = .169, cohen’s d = 0.370, monetary: t(56)=1.1, p = .272, Cohen’s d = 0.292).

To explore whether the time spent on training affected anticipatory reward processing in the midbrain, we ran a *post hoc* analysis with total time spent on training for each participant as a covariate. Adding this covariate (and interaction terms) did not change the results (Reward x Domain x Time x Intervention: F(1,55)=7.4, p = .009, η_p_^2^ = 0.118). Because BMI and waist showed effects of the intervention (see secondary Results: Anthropometric measures), we added BMI and waist as covariates to the analysis. This also did not change the results qualitatively (Reward x Domain x Time x Intervention interaction with BMI covariate: F(1,55)=5.93, p=.018, η_p_^2^ = 0.097; with waist circumference covariate: F(1,55)=5.94, p=.018, η_p_^2^ = 0.097).

#### Reward Receipt

The intervention did not affect neural responses during the receipt of reward. Specifically, no significant main effects of Intervention or interactions with Intervention were found for BOLD responses to reward receipt in whole-brain analyses, nor in ROI analyses using *a priori* defined ROIs for striatum and midbrain. For main effects and other interaction effects of reward receipt see **Table 4**.

**Table 4.** Reward Receipt. Summary of brain regions exhibiting main effects of reward, domain and/or interactions with domain, training, and time.

| Label | Side<br>(Left/Right) | MNI-coordinates<br>x, y, z (mm) |  |  | Size (number of<br>voxels) | *p*FWE<br>(peak-level) | *t*-value <sup>a</sup><br>(peak) |
| --- | --- | --- | --- | --- | --- | --- | --- |
| Main effect of receipt: hits (high > low) > too lates (high > low)<sup>b</sup> |  |  |  |  |  |  |  |
| Nucleus accumbens | L | -14 | 6 | -12 | 880 | < .001 | 13.61 |
| Putamen | L | -18 | 10 | -6 |  | < .001 | 12.41 |
| Hippocampus | L | -16 | -6 | -16 |  | < .001 | 6.89 |
| Putamen | R | 18 | 8 | -8 | 1019 | < .001 | 12.50 |
|  |  | 22 | 14 | -4 |  | < .001 | 12.42 |
|  |  | 30 | -10 | 2 |  | < .001 | 8.63 |
| Middle temporal gyrus | R | 48 | -72 | 0 | 1371 | < .001 | 9.85 |
| Inferior temporal gyrus | R | 52 | -54 | -16 |  | < .001 | 8.91 |
| Middle occipital lobe | R | 32 | -80 | 10 |  | < .001 | 7.40 |
| Superior frontal gyrus, medial orbital | L | -4 | 50 | -6 | 1197 | < .001 | 9.63 |
|  |  | -6 | 42 | -8 |  | < .001 | 8.35 |
|  |  | -6 | 60 | 2 |  | < .001 | 7.87 |
| Superior frontal gyrus | L | -20 | 30 | 52 | 794 | < .001 | 9.38 |
| Middle frontal gyrus | L | -22 | 18 | 46 |  | < .001 | 6.82 |
| Superior frontal gyrus | L | -14 | 46 | 38 |  | < .001 | 6.07 |
| Inferior temporal gyrus | L | -52 | -48 | -14 | 499 | < .001 | 9.26 |
| Inferior parietal gyrus | L | -48 | -40 | 48 | 1530 | < .001 | 8.86 |
| Superior parietal gyrus | L | -30 | -66 | 48 |  | < .001 | 8.28 |
| Inferior parietal gyrus | L | -42 | -40 | 40 |  | < .001 | 8.14 |
| Inferior parietal gyrus | R | 34 | -48 | 50 | 1074 | < .001 | 8.72 |
| Supramarginal gyrus | R | 46 | -36 | 46 |  | < .001 | 8.69 |
| Inferior parietal gyrus | R | 40 | -42 | 50 |  | < .001 | 7.22 |
| Inferior frontal gyrus, triangular | L | -40 | 36 | 14 | 324 | < .001 | 8.60 |
| Putamen | L | -30 | -12 | 4 | 104 | < .001 | 8.19 |
| Inferior frontal gyrus, triangular | L | -36 | 36 | 12 | 142 | < .001 | 7.12 |
| Inferior frontal gyrus, orbital | L | -26 | 30 | -18 |  | < .001 | 6.70 |
| Caudate nucleus | L | -20 | -8 | 26 | 92 | < .001 | 7.01 |
|  |  | -20 | -16 | 30 |  | < .001 | 6.10 |
| Inferior frontal gyrus, opercular | L | -44 | 6 | 26 | 51 | < .001 | 6.41 |
| Superior frontal gyrus | R | 22 | 30 | 48 | 28 | < .001 | 6.26 |
| Caudate nucleus | R | 18 | -8 | 26 | 43 | .001 | 6.03 |
|  |  | 20 | 6 | 20 |  | .005 | 5.62 |
| Inferior frontal gyrus, orbital | R | 28 | 36 | -14 | 16 | .001 | 5.97 |
| Cingulate gyrus, mid part | R | 4 | -38 | 34 | 27 | .001 | 5.85 |
| Middle occipital lobe | L | -24 | -92 | 8 | 48 | .002 | 5.77 |
|  |  | -32 | -88 | 10 |  | .007 | 5.53 |
| Precentral gyrus | R | 48 | 4 | 28 | 31 | .002 | 5.76 |
| Inferior occipital lobe | L | -48 | -74 | -2 | 12 | .002 | 5.74 |
| Middle occipital lobe | L | -26 | -84 | 20 | 14 | .002 | 5.67 |
| Precuneus | R | 2 | -62 | 24 | 23 | .004 | 5.66 |
| Paracentral lobule | L | -2 | -26 | 60 | 28 | .004 | 5.63 |
|  | R | 8 | -24 | 62 |  | .004 | 5.45 |
| Hippocampus | L | -32 | -34 | -6 | 1 | .007 | 5.55 |
| Inferior parietal gyrus | L | -26 | -50 | 44 | 1 | .035 | 5.19 |
| Middle temporal gyrus | L | -62 | -10 | -22 | 1 | .045 | 5.14 |
| Main effect of receipt: too late (high > low reward) > hits (high > low reward) <sup>b</sup> |  |  |  |  |  |  |  |
| Middle temporal gyrus | R | 48 | -26 | -6 | 1877 | < .001 | 13.37 |
|  |  | 48 | -36 | 0 |  | < .001 | 10.35 |
| Supramarginal gyrus | R | 60 | -42 | 36 |  | < .001 | 9.06 |
| Inferior frontal gyrus, orbital | R | 48 | 22 | -4 | 956 | < .001 | 9.42 |
| Inferior frontal gyrus, triangular | R | 54 | 22 | 4 |  | < .001 | 8.04 |
|  |  | 44 | 22 | 8 |  | < .001 | 7.15 |
| Supramarginal gyrus | L | -62 | -44 | 26 | 342 | < .001 | 8.91 |
| Supplementary motor area | R | 6 | 24 | 62 | 1584 | < .001 | 8.58 |
|  |  | 8 | 14 | 66 |  | < .001 | 8.45 |
| Superior frontal gyrus, medial | R | 4 | 34 | 54 |  | < .001 | 7.70 |
| Middle temporal gyrus | L | -50 | -28 | -4 | 437 | < .001 | 8.33 |
|  |  | -50 | -48 | 8 |  | < .001 | 6.95 |
| Thalamus | L | -8 | -16 | 8 | 355 | < .001 | 7.51 |
| Thalamus | R | 10 | -16 | 10 |  | < .001 | 7.35 |
|  |  | 8 | -8 | 6 |  | < .001 | 7.02 |
| Middle frontal gyrus | R | 30 | 50 | 24 | 274 | < .001 | 7.50 |
| Middle temporal gyrus | L | -56 | 2 | -14 | 626 | < .001 | 7.29 |
| Insula | L | -34 | 22 | -8 |  | < .001 | 7.14 |
|  |  | -36 | 20 | 8 |  | < .001 | 6.80 |
| Postcentral gyrus | L | -40 | -22 | 50 | 320 | < .001 | 7.04 |
|  |  | -48 | -20 | 46 |  | < .001 | 6.30 |
|  |  | -42 | -24 | 38 |  | .033 | 5.21 |
| Middle frontal gyrus | L | -26 | 48 | 24 | 90 | < .001 | 6.89 |
|  |  | -24 | 38 | 20 |  | .013 | 5.41 |
| Caudate nucleus | R | 12 | 2 | 14 | 20 | < .001 | 6.24 |
| Cingulate gyrus, mid part | R | 6 | -18 | 36 | 15 | < .001 | 6.05 |
| Middle temporal pole | R | 52 | 8 | -22 | 32 | .001 | 5.92 |
| Insula | R | 40 | 0 | -14 | 13 | .002 | 5.80 |
| Caudate nucleus | L | -8 | 8 | 6 | 4 | .003 | 5.71 |
| Cerebellum | R | 22 | -50 | -22 | 7 | .004 | 5.66 |
| Rolandic operculum | L | -40 | -20 | 18 | 9 | .009 | 5.48 |
| Superior temporal gyrus | L | -40 | -4 | -14 | 2 | .016 | 5.37 |
| Middle temporal gyrus | R | 50 | 2 | -22 | 1 | .038 | 5.17 |
| Temporal pole: superior temporal gyrus | R | 50 | 16 | -18 | 1 | .044 | 5.14 |
| Midbrain | R | 6 | -24 | -6 | 1 | .046 | 5.13 |
| Calcarine fissure | R | 8 | -80 | 12 | 1 | .047 | 5.13 |
| Interaction effect of domain x reward: caloric (hits (high > low reward) > toolates (high > low reward)) > monetary (hits (high > low reward) > toolates (high > low reward))<sup>b</sup> |  |  |  |  |  |  |  |
| Lingual gyrus | R | 18 | -86 | -4 | 195 | < .001 | 8.96 |
| Calcarine fissure | L | -10 | -90 | -4 | 63 | .001 | 6.04 |
| Cerebellum | R | 16 | -80 | -16 | 5 | .005 | 5.59 |
| Interaction effect of Intervention x Domain x Reward<sup>b</sup> |  |  |  |  |  |  |  |
| Putamen | R | 30 | -14 | 8 | 1 | .015 | 5.38 |
| Interaction effect of domain x receipt x reward: monetary (hits (high > low reward) > toolates (high > low reward)) > caloric (hits (high > low reward) > toolates (high > low reward))<sup>b</sup> |  |  |  |  |  |  |  |
| Lingual | R | 18 | -84 | -4 | 116 | < .001 | 7.02 |
| Lingual | L | -14 | -88 | -4 | 1 | .023 | 5.01 |

<sup>a</sup> Degrees of freedom: 1,224; <sup>b</sup> $p < .05$ , whole-brain FWE corrected.

### Anthropometric outcomes

As a secondary outcome, we analyzed the effects of the two interventions on the anthropometric measures. Although we initially intended to assess only BMI and waist-to-hip ratio (WHR), we have added the analysis of waist circumference as an additional exploratory measure to assess abdominal obesity^49^. Changes in abdominal obesity as measured with WHR may be masked because of the relative nature of the measure (i.e., if the interventions affect waist and hip circumference similarly, especially in women^50^). The interventions had differential effects on the anthropometric measures as indicated by a significant Time x Intervention interaction. Specifically, the active control, EC, intervention resulted in both decreased BMI and waist circumference (main Time: BMI: F(1,25)=6.2, p=.020, η_p_^2^ = 0.198; waist circumference: F(1,25)=17.9, p<.001, η_p_^2^ = 0.418), whereas the ME intervention did not affect either of them (main Time: BMI: F(1,31)<1, p=.648, η_p_^2^ = 0.007; waist circumference: F(1,31)<1, p=.504, η_p_^2^ = 0.015). Waist-to-hip ratio was not affected by either of the interventions (Time x Intervention: F(1,56)<1, p = .379, η_p_^2^ = 0.014). For all comparisons see **Table 5**.

**Table 5.**
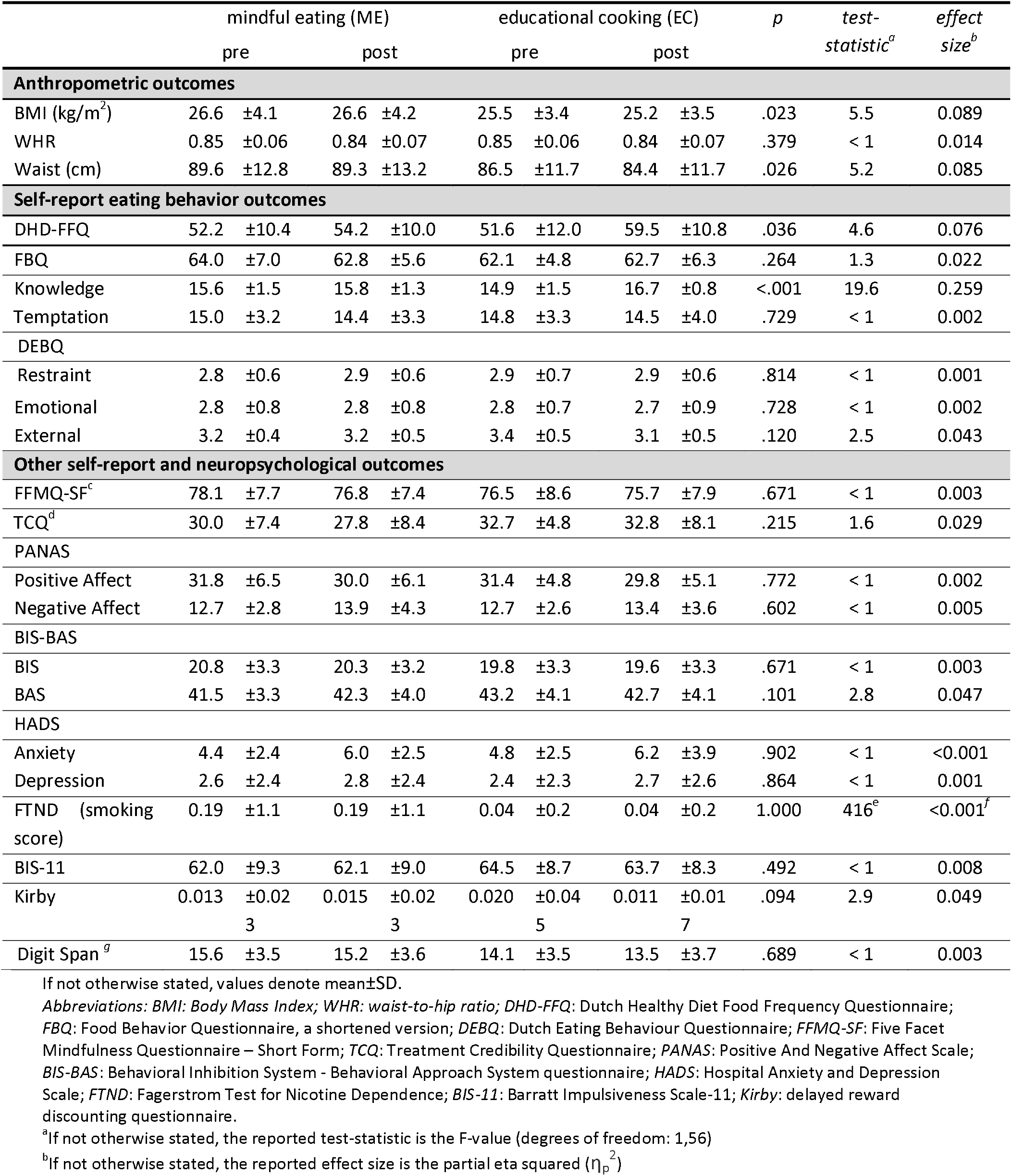
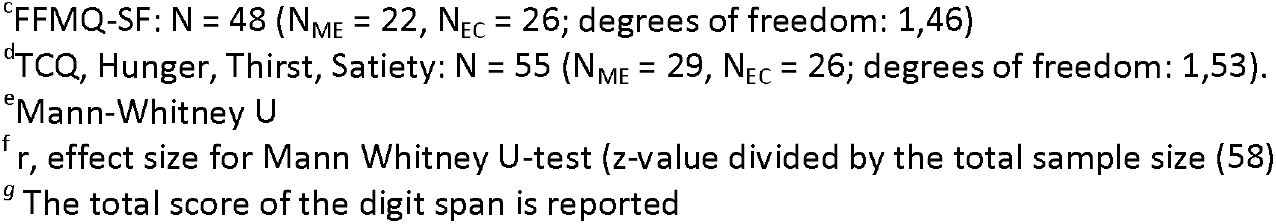
Secondary anthropometric, self-reported eating behaviour, and neuropsychological outcomes. Means and standard deviations, pre- and post-training, for each group (mindful eating, ME; educational cooking, EC) separately, and Time (pre, post) x Intervention (ME, EC) statistics.

### Self-reported and neuropsychological outcomes

As another secondary outcome, we assessed intervention effects on eating-related self-reported measures. We found that EC participants reported closer compliance to the Dutch food-based guidelines for healthy eating (main Time: F(1,25)=12.8, p=.001, η_p_^2^ = 0.339) than ME participants following their intervention (main Time: F(1,31)=1.4, p=.244, η_p_^2^ = 0.044), as substantiated by a significant Time x Intervention interaction for DHD-FFQ scores. EC participants also showed a significant increase in knowledge on healthy eating following the intervention (main Time: F(1,25)=48.8, p<.001, η_p_^2^ = 0.661), whereas ME participants did not (main Time: F(1,31)<1, p=.394, η_p_^2^ = 0.024), as evidenced by a significant Time x Intervention interaction for FBQ scores. The other sub-scale of the FBQ (temptation) did not show any differential intervention effects; neither did any of the sub-scales of the DEBQ (restraint, emotional, and external eating). For all comparisons see **Table 5**.

Analysis of the other self-reported and neuropsychological measurements – including those related to the intervention (FFMQ-SF, TCQ), affect (PANAS, BIS-BAS, HADS), impulsivity (FTND, BIS-11, Kirby), and working memory (digit span) revealed no significant interactions between Time and Intervention (**Table 5**).

To establish whether the observed differential intervention effects (Time x Intervention interactions) in the anthropometric and eating-related self-report measures were long-lasting, we ran *post hoc* analyses by adding the one-year follow-up data as an extra level of factor Time (pre, post, follow-up) in the ANOVAs for all participants from the reported sample that returned for the follow-up (ME: n=26, EC: n=20)(**Figure 4**). For BMI and waist circumference, degrees of freedom were corrected using Huynh-Feldt estimates of sphericity due to violation of the sphericity assumption. BMI, WHR, and waist circumference did not show any long-term intervention-related changes (Intervention x Time, BMI: F(1.550,69.77)<1, p=0.468, η_p_^2^ = 0.015; WHR: F(2,44)<1, p=0.589, η_p_^2^ = 0.024; waist: F(1.742,78.384)=2.213, p=0.123, η_p_^2^ = 0.047). BMI, WHR, and waist circumference changed over time irrespective of the intervention (main Time, BMI: F(1.550,69.77)=3.730, p=0.039, η_p_^2^ = 0.077; WHR: F(2,44)=5.099, p=0.010, η_p_^2^ = 0.188; waist: F(1.742,78.384)=4.837, p=0.014, η_p_^2^ = 0.097). Planned *post hoc* comparisons revealed that the non-significant intervention effects on the anthropometric measures – after including the follow-up time point – were caused by a lack of significant differences between pre-intervention measurements and one-year follow-up measurements for either group (all p>0.1). This means that the BMI- and waist circumference-reducing effects of the active control (EC) intervention (versus the mindful eating intervention) were no longer visible at one-year follow-up.

**Figure 4.**
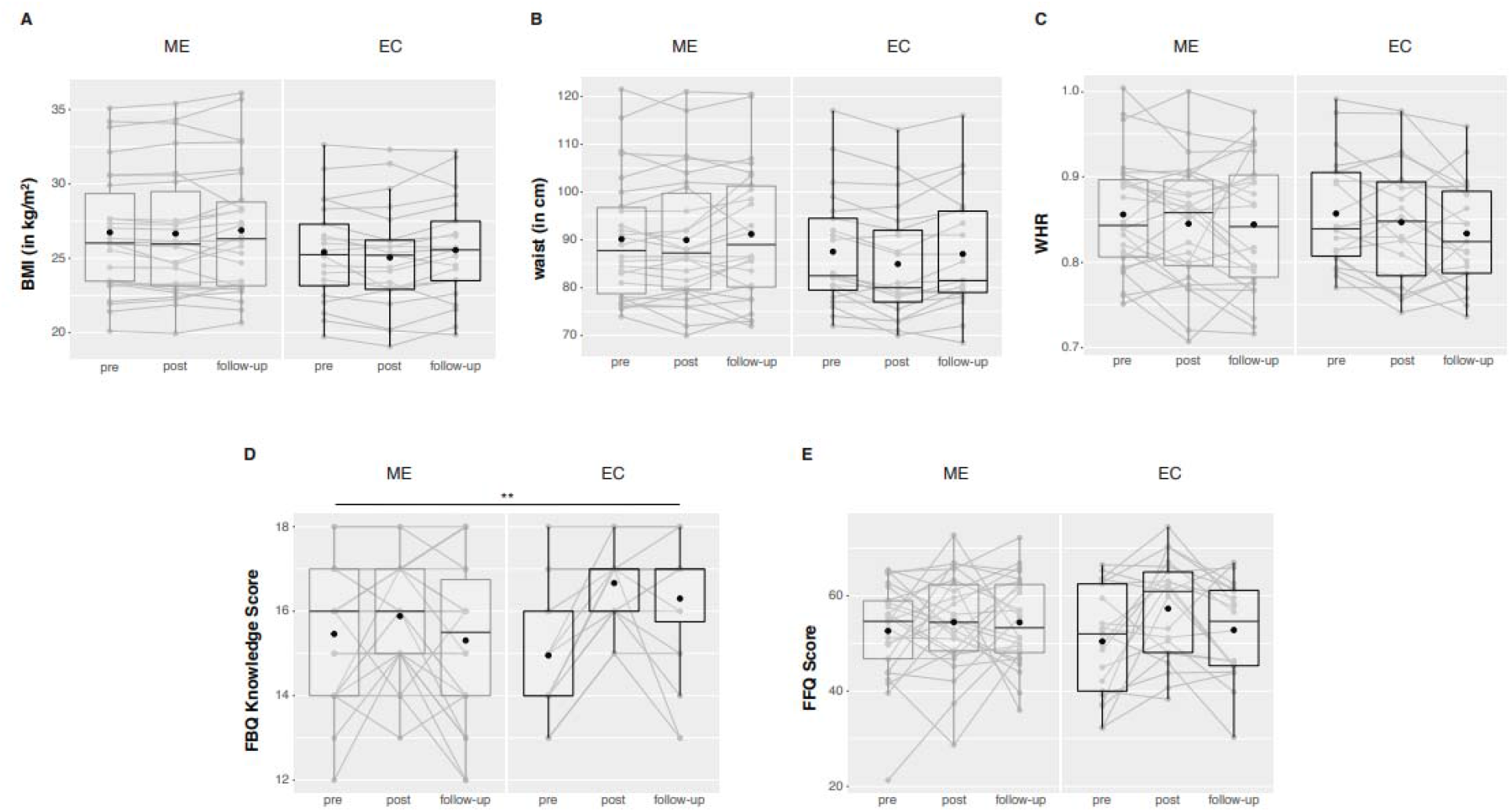
Anthropometric (upper panels) and eating-related self-report measures (lower panels) 1 year after the intervention. No long-lasting intervention effects were observed for (A) BMI, (B) waist circumference, (C) waist-to-hip ratio (WHR), and (E) compliance to the Dutch guidelines for healthy diet (DHD-FFQ). Only knowledge on healthy eating (D) remained high following the educational cooking (EC) intervention. No intervention effects were observed for the mindful eating (ME) group. Box plots show the median and interquartile range, with the black dot denoting the mean. Note that the medians of the EC group in figure (D) do not fall in the interquartile range. Individual data points at the different test sessions are connected for illustrative purpose. ** asterisks denote a significant Time x Intervention interaction with p < 0.01.

The EC-related increase in knowledge on healthy eating remained significant after including the one-year follow-up time point (Intervention x Time, FBQ knowledge: F(2,44)=7.4, p=0.002, η_p_^2^ = 0.253), caused by a lingering increase in knowledge on healthy eating for EC participants at one-year follow-up relative to pre-intervention measurements (F(1,20)=17.06, p=0.001, η_p_^2^ = 0.460). In contrast, the effects of the active control (EC) intervention on self-reported compliance to the Dutch guidelines for healthy diet were not long lasting (Intervention x Time, DHD-FFQ: F(2,43)=2.121, p=0.132, η_p_^2^ = 0.090). Similar to the anthropometric measures, planned post-hoc comparisons revealed that these DHD-FFQ scores were comparable between the pre-intervention and one-year follow-up measures for either group (all p>0.1), meaning that the previously observed post-pre effects of the active control (EC) intervention were only short lasting.

## Discussion

The primary objective of this study was to investigate the effects of an 8-week mindful eating intervention on striatal reward anticipation responses as well as response times during an incentive delay task. In addition to the striatum, we explored these effects in the midbrain – as part of the mesolimbic reward circuit with its dopaminergic projections to the striatum^4,5^ – as regions of interests (ROIs). We observed that mindful eating training significantly impacted reward anticipation in the midbrain relative to the active control training, with relatively reduced caloric versus monetary reward responses in this region after the intervention. We found no effect of the interventions in the striatum or on response times during the incentive delay task. Anthropometric measures of obesity (i.e. secondary outcome: BMI) temporarily decreased and self-reported (knowledge of) healthy food intake (i.e. secondary outcome: eating behavior questionnaires) increased following the educational cooking intervention, but not following the mindful eating intervention.

We did not observe any intervention effect on the response times or on striatal fMRI (BOLD) responses during the incentive delay task. Previous studies have shown that greater subcortical reward responses to caloric cues, particularly in striatum, are associated with obesity ^51,52^, with weight gain ^9^, and with increased snack food intake in healthy-weight to overweight individuals ^7^. Despite this clear involvement of striatum in food reward anticipation and its relationship with eating behavior, we found no effects of mindful eating training on striatal BOLD responses. Below, we interpret these null results.

We did however observe intervention effects on midbrain reward anticipation in the current study, with relatively reduced responses to the caloric (i.e. high-calorie drink versus water) compared with the monetary (50 ct. versus 1 ct.) cues – in an, on average, overweight sample of participants motivated to improve their dietary habits. Dopaminergic midbrain neurons are crucial for processing predicted reward value ^5,53^ and, in concert with striatum, modulate motivated behavior such as eating ^54^. In line with this, Small et al. ^55^ showed that midbrain activity, as measured with positron emission tomography (H_2_^15^O), decreased with reduced self-reported reward value of chocolate in a sample of healthy individuals consuming chocolate beyond satiety. In another study, midbrain BOLD responses to sips of palatable milkshake were found to positively correlate with subsequent *ad libitum* milkshake intake in a group of healthy-weight to moderately obese individuals ^56^. Moreover, overweight and obese compared with normal weight adolescents showed increased activations in midbrain during anticipation of decisions involving risk and reward ^12^. Furthermore, both midbrain and striatal BOLD responses to palatable food pictures were found to correlate positively with self-reported reward drive in healthy individuals ^57^. These (indirect) measures of motivated eating behavior are thus associated with greater mesolimbic responses when processing food reward value.

Our finding that anticipatory midbrain responses were relatively reduced in the caloric versus monetary domain is in line with a previous study showing that only a brief 50-min mindful eating workshop (versus an educational video) reduced subsequent impulsive choice patterns for food-, but not money-related outcomes ^58^. However, in studies comparing meditators with non-meditating controls, meditators exhibited reduced striatal BOLD responses to primary reward prediction errors ^25^ as well as monetary reward anticipation ^24^. In the latter study, Kirk and colleagues ^24^ compared meditators to non-meditators without a baseline measurement. The observed decrease in striatal reward processing could thus be due to pre-existing between-group differences ^59^. Since the present study was actively controlled including pre and post measurements, the current effects can be more reliably ascribed to the mindfulness intervention. Kirk and colleagues ^26^ also performed a similar randomized actively controlled study including pre and post measurements and found that vmPFC value signals were modulated by the mindfulness intervention for both primary (juice) and secondary (monetary) rewards. These general reward effects versus our relative caloric versus monetary effects might be due to both the type of intervention (general MBSR in Kirk et al. ^26^ versus mindful eating presently) as well as the study sample. Specifically, in our study, participants were highly motivated to change undesired eating habits and their mindfulness practice was targeted at overcoming those – including homework practices such as resisting impulsive eating behaviors.

Moreover, note that we did not observe any effects of either the ME or the EC intervention on neural responses at the time of caloric or monetary reward receipt. One might have expected reductions in vmPFC BOLD responses following the mindfulness-based intervention as was reported by Kirk et al. ^26^ for juice delivery. However, another important difference with the current study is that we used promised (i.e. delivered after scanning) instead of actual rewards (delivered during scanning). Moreover, our design was optimized for reward anticipation, with perhaps not enough successful reward receipt trials (i.e. approximately 33% of all anticipated rewards were missed). Together, our results suggest that a targeted mindful eating – instead of general mindfulness – intervention may have more specific effects on caloric versus monetary reward anticipation.

The specificity of our results for midbrain, not striatum, finds resonance in a study in healthy individuals by O’Doherty and colleagues ^60^, who found significant responses to cues predicting the receipt of a glucose solution versus a neutral taste in midbrain only, whereas both midbrain and striatum were responsive to cues predicting the receipt of a sweet versus an aversive salty taste. The latter contrast may be a larger one in terms of valence, which might implicate that our caloric versus water contrast was not sensitive enough to show intervention effects in the striatum – despite showing main task effects of reward anticipation. Given the coding of predicted reward in the midbrain, we speculate that the currently observed relative effect of the mindful eating intervention on anticipatory midbrain responses to caloric versus monetary cues suggests that mindful eating practice may be able to reduce the impact of food cues on reward processing.

The question then arises whether mindfulness affects midbrain responses through top-down or bottom-up processes. Current theories on mindfulness-based interventions emphasize that improvements in emotion regulation occur through increased prefrontal cortex-mediated top-down control of regions processing affect, such as the amygdala ^61,62^. An alternative way to reducing incentive motivation is through extinction during mindfulness practice, akin to exposure therapy ^61,62^, which would rather be a bottom-up process. Practicing mindful eating requires one to actively withhold or interrupt cue-triggered eating, a process that may lead to extinction of conditioned responses to highly caloric stimuli ^61,63,64^ as well as the formation of new memories related to those stimuli (i.e., not reacting to them). As a result, choices for high caloric foods may be further reduced ^65,66^. However, incentive motivation could also be reduced through other bottom-up effects on, for example, physiological state rather than through extinction. Increased awareness of states like hunger or satiety ^67^ are known to modulate conditioned responses to reward-related cues ^68^. Future confirmatory studies are needed to verify the exploratory midbrain findings and investigate the underlying bottom-up versus top-down mechanisms, for instance by employing tasks manipulating top-down control on food reward processes, addressing the effects of physiological state and interoception, and by employing connectivity analyses between cortical and mesolimbic regions.

The present mindful eating effects on caloric versus monetary reward anticipation in the midbrain were not accompanied by changes in our secondary outcome measures related to real-life eating behavior, i.e., reductions in weight, waist-hip ratio or waist circumference, or changes in self-reported eating behavior. Several other studies have found that an intensive mindful eating intervention did lead to reduced measures associated with overeating such as consumption of sweets ^22^, binges, externally and emotionally driven eating ^69^ and reductions in BMI ^20^ in non-clinical populations, as well as number of binges in binge-eating disorder ^18^. On the other hand, a more recent review by Warren and colleagues concludes that there is a lack of compelling evidence of mindfulness and mindful eating interventions leading to a reduction in weight ^70^. The current lack of mindful eating intervention-related reductions in our secondary measures of (abdominal) obesity might reflect the heterogeneity of our sample, including normal-weight, overweight and obese individuals; with larger mindfulness-related reductions in food intake seen in overweight and obese populations in previous studies ^70^. Moreover, the study design – including sample size – was optimized for the primary outcome measure (i.e., neural effects) and plausibly less optimal for showing these behavioral effects after the mindful eating intervention. We were also not able to show increased self-reported mindfulness after the intervention on the established short version of the Five Facet Mindfulness Questionnaire^31^, but this questionnaire was only employed in a sub group (in n=22 of the total n=32 ME vs n=26 EC). In fact, ineffectiveness of our mindful eating intervention is highly unlikely given the observed midbrain findings in the hypothesized direction here (although exploratory) and our previously published effects on behavioral flexibility ^71^. Sampling a greater and more homogeneous population in terms of BMI is advised for future studies to be able to demonstrate a link between reduced mesolimbic reward responses and altered eating behavior following a mindful eating intervention. For now, it is unclear how mindfulness-induced reductions in midbrain responses to caloric versus monetary reward anticipation contribute to changes in real-life eating behavior.

In contrast, we did observe beneficial effects of the educational cooking intervention on anthropometric measures of obesity and self-reported eating behavior, whereas this group did not demonstrate any intervention effects on mesolimbic reward anticipatory responses. The beneficial effects might not be surprising for this group, since the educational cooking intervention was explicitly aimed at promoting healthy food intake, with reduced intake of sugar, fats and salt as part of the homework assignments. This led to short-term reductions in weight and waist circumference, as well as increased self-reported adherence to the Dutch healthy diet (DHD-FFQ). Given those health benefits of the educational cooking intervention and the relatively reduced food reward anticipation responses of the mindful eating intervention, it might be fruitful to develop a combined program for therapeutic practice or for preventive strategies. Although weight control and diet interventions are often successful in producing significant weight loss on the short term, they often fail to produce long-term weight maintenance ^72^. This is supported by our analyses of BMI, waist circumference, and self-reported compliance to the Dutch healthy diet guidelines (DHD-FFQ) at one-year follow-up in the present study. These secondary measures returned to baseline one year after the educational cooking intervention, despite the fact that knowledge of healthy eating remained significantly higher compared with baseline in the educational cooking group. Previous studies investigating factors contributing to successful weight maintenance have shown that reductions in subcortical responses to food reward cues may be beneficial for prevention or treatment of obesity ^7–9^. Therefore, we speculate that a combination of the two interventions with a focus on both information and behavior might lead to longer-lasting health benefits than either intervention on its own.

We note that the lack of a – likely, very subtle - effect of mindful eating, e.g. on striatal reward anticipation, might well reflect the inclusion of a well-matched active control intervention; enabling the observed effects to be actually attributed to mindfulness practice ^62^. The randomized, active-controlled nature of the study was probably also the reason for a high dropout rate. This may reflect a lack of motivation to take part in the interventions, although the dropout rate in the mindful eating group was more than 15% lower (non-significantly) than in the active control condition (that showed clear effects on secondary outcome measures). We speculate that dropout rates could have been lower and motivation higher in the current study had we been able to offer participants to take part in the other intervention program after completion of the study, which is commonly done for mindfulness studies that include a waitlist control group. To address differences in results of previous mindfulness or meditation studies without active control condition, future mindfulness intervention studies, especially those aimed at unraveling subtle mechanistic effects, are recommended to not only include a well-matched active control intervention but also a waitlist control group.

In conclusion, we found that an intensive mindful eating intervention reduced midbrain food, relative to monetary, reward anticipation. These results have to be confirmed in future studies, as we primarily hypothesized striatal effects, and the midbrain findings are the result of exploratory analyses. Future studies are also required to demonstrate the clinical relevance of mindfulness-mediated reductions in food anticipation for counteracting reward cue-driven overeating, particularly given that we did not observe mindfulness-related changes in anthropometric or eating behavior measures. Given the success of mindfulness-based programs in reducing symptoms of other reward-related disorders such as substance use ^73,74^ and problem gambling ^75^, our findings of relatively specific reduced anticipatory reward responses may also be relevant for these other targets of abuse.

## Acknowledgements

This work was supported by a VENI grant (016.135.023) of The Netherlands Organization for Scientific Research (NWO) and an AXA Research Fund fellowship (Ref: 2011) to E.A. R.C. was supported by the James S. McDonnell Foundation (220020328) and a VICI grant (453-14-005) of NWO.

The authors wish to thank the participants and all people who were involved in developing, teaching, and arranging the logistics of the intervention programs. Specifically, we would like to thank Ellen Jansen and Nicole Schoonbrood of the Radboud university medical center for Mindfulness for developing and teaching the mindful eating intervention. We would also like to thank Desiree Lucassen for developing the educational cooking intervention in collaboration with the Division of Human Nutrition and Health of Wageningen University, and for teaching it. She was assisted by chef Pieter Paul de Grood and bachelor student Hanne de Jong. In addition, we are thankful to Susanne Leij-Halfwerk and Suzan de Bruijn of the department of Nutrition and Dietetics at HAN University of Applied Sciences for access to their cooking classrooms. Furthermore, the authors are grateful to Marcel Zwiers for helpful comments regarding ICA AROMA, and to Vishnu Murty and Ian Ballard for advice on the use of a probabilistic midbrain ROI.

## Author contributions

EA acquired funding for the study. EA, LKJ, RC, AEMS, and JHMdV designed the study. LKJ, ID, IvL, and JW acquired and analyzed the data, supervised by EA and RC. AEMS and JHMdV supervised the execution of the interventions. LKJ, ID, and EA wrote the first version of the manuscript. All authors corrected the manuscript and approved it for final submission.

## Data availability

The datasets analysed during the current study are available on http://dx.doi.org/10.17632/fthcv3kns9.1.

